# Machine learning-based tissue of origin classification for cancer of unknown primary diagnostics using genome-wide mutation features

**DOI:** 10.1101/2021.10.05.463244

**Authors:** Luan Nguyen, Arne van Hoeck, Edwin Cuppen

**Author notes:** shared last authors (Arne van Hoeck, Edwin Cuppen).

## Abstract

Tumor tissue of origin (TOO) is an important factor for guiding treatment decisions. However, TOO cannot be determined for ~3% of metastatic cancer patients and are categorized as cancers of unknown primary (CUP). As whole genome sequencing (WGS) of tumors is now transitioning from the research domain to diagnostic practice in order to address the increasing demand for biomarker detection, its use for detection of TOO in routine diagnostics also starts becoming within reach. While proof of concept for the use of genome-wide features has been demonstrated before, more complex WGS mutation features, including structural variant (SV) driver and passenger events, have never been integrated into TOO-classifiers even though they bear highly characteristic links with tumor TOO. Using a uniformly processed dataset containing 6820 whole-genome sequenced primary and metastatic tumors, we have developed Cancer of Unknown Primary Location Resolver (CUPLR), a random forest based TOO classifier that employs 502 features based on simple and complex somatic driver and passenger mutations. Our model is able to distinguish 33 cancer (sub)types with an overall accuracy of 91% and 89% based on cross-validation (n=6139) and hold out set (n=681) predictions respectively. We found that SV derived features increase the accuracy and utility of TOO classification for specific cancer types. To ensure that predictions are human-interpretable and suited for use in routine diagnostics, CUPLR reports the top contributing features and their values compared to cohort averages. The comprehensive output of CUPLR is complementary to existing histopathological procedures and may thus improve diagnostics for patients with CUP.

## Introduction

Cancers of unknown primary (CUPs) is an umbrella term for advanced stage metastatic tumors for which the tumor tissue of origin (TOO) cannot be conclusively determined based on routine diagnostics (typically via histopathology [1]), and there is also a significant fraction of patients with indeterminate or differential diagnoses, especially with poorly differentiated tumors [2]. Patients with uncertain TOO diagnosis suffer from a lack of therapeutic options as primary cancer type classification is a dominant factor in guiding treatment decisions [3].

Thus far, the most performant TOO classifiers have been developed on data from whole genome sequencing (WGS) [4,5], whole transcriptome sequencing (WTS) [6,7] or methylation profiling [8]. While gene expression and methylation profiling for cancer diagnostics is not yet widespread [9,10], WGS is being increasingly adopted, driven by its utility for comprehensive identification of actionable mutations [11]. WGS-based TOO classifiers are thus at this moment more viable for application in the clinic. Recently developed WGS-based classifiers [4,5] were shown to outperform targeted or whole exome sequencing based approaches [12,13] due to being able to utilize mutations from all genomic regions. The main features employed by these classifiers included mutational signatures which are patterns of somatic mutations resulting from exogenous or endogenous mutational processes (e.g. C>T mutations due to ultraviolet radiation exposure in melanoma) [14], as well as regional mutational density (RMD) which represents the genomic distribution of somatic mutations that are associated with tissue type specific chromatin states, whereby late-replicating closed chromatin regions show increased mutation rates [15]. However, not all WGS based features are yet fully explored for TOO classification including complex mutagenic features such as viral DNA integrations, driver gene fusions, and other complex structural events (e.g. chromothripsis), as well as non-mutagenic features such as gender, all of which have been shown to be correlated with specific tumor type(s). Indeed, human papillomavirus (HPV) sequence insertions are specifically and frequently found in cervical and head and neck cancer [14], *KIAA1549-BRAF* fusions in pilocytic astrocytomas [13], and liposarcomas frequently harbor *FUS-DDIT3* fusions [15] and chromothripsis events [16].

Here we describe the development of CUPLR (<u>C</u>ancer of <u>U</u>nknown <u>P</u>rimary <u>L</u>ocation <u>R</u>esolver), a TOO classifier that integrates current state-of-the-art WGS based mutation features as well as thus far unexplored complex structural variant (SV) features. CUPLR comprises an ensemble of binary random forest classifiers that each discriminate one of 33 cancer types with an overall accuracy of 0.91. We find that while RMD and mutational signatures were highly predictive of cancer type (in line with existing classifiers [4,5]), the incorporation of SV features improves prediction performance for cancer types that currently lack highly informative features. Furthermore, we have ensured that the output of CUPLR, namely the prediction probabilities and the features supporting each prediction, are human interpretable to facilitate diagnostic use and clinical decision making with CUPLR.

## Results

### Extraction of genomic features

To develop CUPLR, we constructed a harmonized dataset from two large pan-cancer WGS datasets from the Hartwig Medical Foundation (Hartwig) and Pan-Cancer Analysis of Whole Genomes consortium (PCAWG). The raw sequencing reads were analyzed with the same mutation calling pipeline to construct a catalogue of uniformly called simple and complex mutations. The harmonized dataset consisted of tumors from 6820 patients across 33 different cancer types (**Figure 1a**). In contrast to many previously published papers [4,5,12,13], this dataset includes a large proportion of samples taken from metastatic lesions, which is relevant for TOO classification as CUP samples are by definition from patients with metastatic cancer.

**Figure 1:**
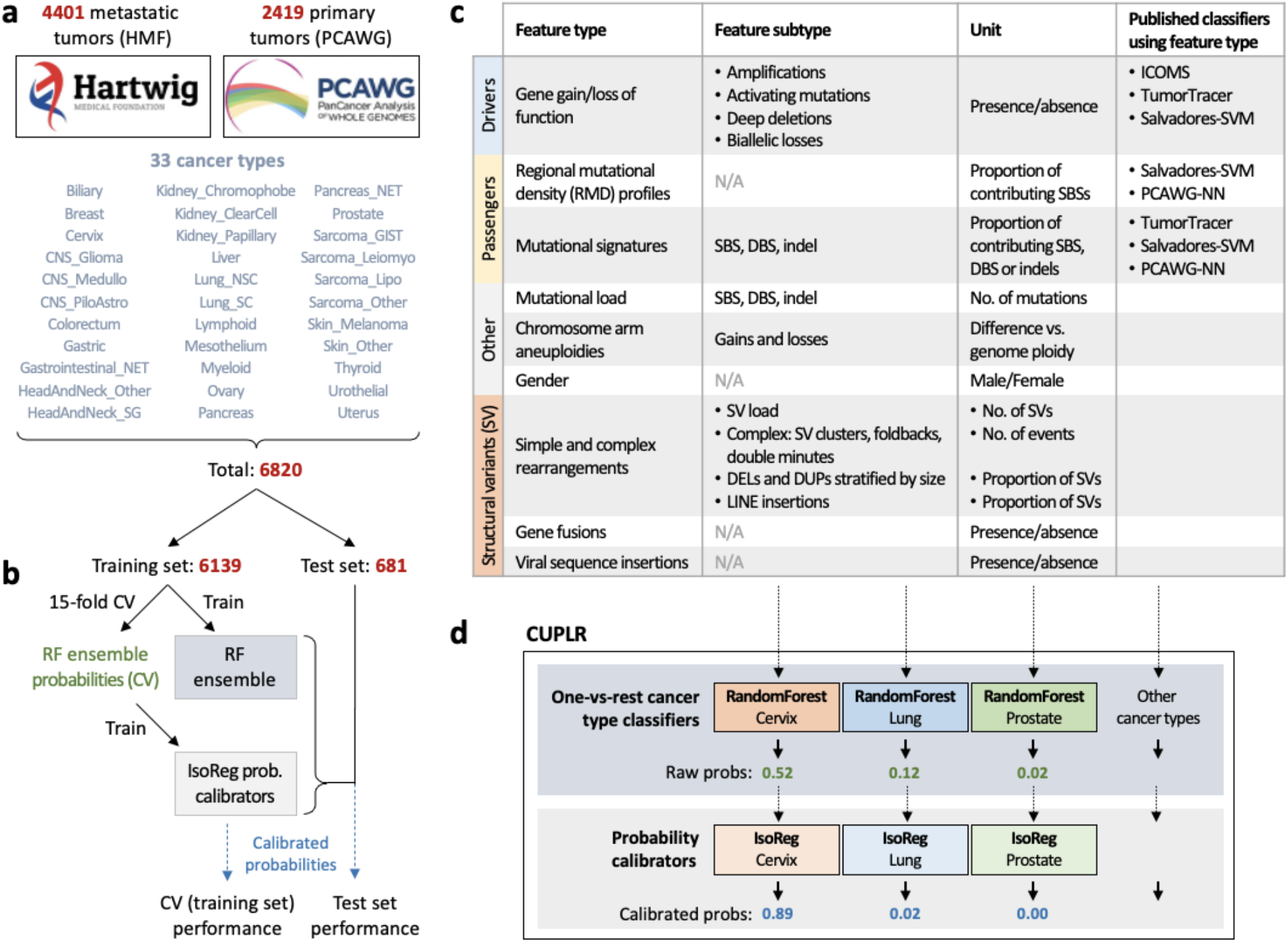
Cancer of Unknown Primary Location Resolver (CUPLR) classifies 33 different cancer types using features derived from all mutation types. **(a)** CUPLR was developed using whole-genome sequencing data 4401 metastatic tumors from the Hartwig Medical Foundation (Hartwig) and 2419 primary tumors from the Pan-Cancer Analysis of Whole Genomes (PCAWG) consortium, totaling 6820 samples across 33 different cancer types. **(b)** 6139 samples were used to train CUPLR. The whole training set was used to train the final random forest ensemble. 15-fold cross-validation was performed to obtain the random forest cancer type probabilities on the training set, which were then used to train the ensemble of isotonic regressions (for probability calibration). CUPLR is composed of the random forest and isotonic regression ensembles as shown in **(d)**. The performance of CUPLR was assessed using the calibrated cross-validation probabilities as well as probabilities obtained by applying CUPLR to the test set. (**c**) A summary of the genomic features extracted from the whole-genome sequencing data and used by CUPLR. The names of the published classifiers refer to the following studies; ICOMS: Inferring Cancer Origins from Mutation Spectra, Dietlen *et al.* 2014 [12]; TumorTracer: Marquard *et al.* 2015 [13]; Salvadores-SVM: support vector machine by Salvadores *et al.* 2019 [5]; PCAWG-NN: PCAWG neural network by Jiao *et al.* 2020 [4]. **Abbreviations**; *RF*: random forest, *IsoReg*: isotonic regression, *CV*: cross-validation, *SBS*: single base substitutions, *DBS*: double base substitutions, *SV*: structural variants, *DEL*: structural deletions, *DUP*: structural duplications, *LINE*: long interspersed nuclear element.

A wide range of features (n=4122) were extracted for classifying cancer type based on driver/passenger and simple/complex mutations (**Figure 1c**). First, we determined the presence of gain of function (amplifications and activating mutations) and loss of function (deep deletions and biallelic loss) events in 205 cancer associated genes. These genes were selected based on having enrichment of gain and/or loss of function events in at least one cancer type (see methods). Second, we calculated the mutational load of single base substitutions (SBS), double base substitutions (DBS), and indels for each sample. Third, we determined the number of contributing mutations to the SBS, DBS and indel signatures from the COSMIC catalog [14]. Fourth, the number of SBSs in each 1Mb bin across the genome (n=3071) was calculated to determine the RMD [17]. Mutational signatures and RMD were normalized by the mutational load of the respective mutation type to account for differences in mutational load across samples. Fifth, for each sample we determined the copy number change of each chromosome arm relative to the genome ploidy [18]. Sixth, gender was determined based on sex chromosome ploidy [19]. Lastly, we parsed the called simple and complex structural variants to determine: (i) the SV load per sample; (ii) the number of large deletions and duplications stratified by size, as well as the number of long interspersed nuclear element (LINE) insertions, normalized by the total number of SVs; (iii) the presence of complex SV events; and (iv) the presence of gene fusions and viral sequence insertions [19,20].

### Classifier training

The extracted genomic features were then used to develop CUPLR, a classifier consisting of two components (**Figure 1d**). Firstly, an ensemble of binary random forest classifiers each discriminates one cancer type versus other cancer types (i.e. one-vs-rest). We chose to use an ensemble of binary classifiers as opposed to one multiclass classifier so that feature selection could be performed per cancer type, since different features are important for each cancer type. Additionally, we chose to use random forests over other algorithms (e.g. neural networks) as they can natively handle different feature types (continuous, Boolean, categorical, etc.) without requiring feature values to be scaled, which also improves model interpretability. The second component of CUPLR is an ensemble of isotonic regressions to calibrate the probabilities produced by each random forest. Random forests tend to be overconfident at probabilities towards 0 and underconfident at probabilities towards 1, and this bias varies between random forests [21]. The calibration we have performed here ensures that probabilities are comparable between random forests. Furthermore, calibration allows for the probabilities to have the following intuitive interpretation: a probability of e.g. 0.8 means that there is an 80% chance of a prediction being correct (this relationship does not hold for raw random forest probabilities).

We used 6139 samples for training and held out 681 samples as an independent test set, with both having the same cancer type proportions. Training of the main random forest ensemble involved several steps. Briefly, due to the sheer number of RMD bins (3071) as well as their sparsity, non-negative matrix factorization (NMF) was for each cancer type performed on the RMD bins to reduce these to 44 cancer type-specific RMD profiles. Then, for each cancer type, univariate feature selection was performed (to remove irrelevant features) with 502 features ultimately being selected (219 numeric and 283 boolean; **Supplementary data 3**). This was followed by class resampling (to alleviate imbalances in the number of samples for each cancer type), and subsequently training of the binary random forest itself (**Supplementary figure 1**). The above training procedure was applied to all samples of the training set to produce the final random forest ensemble. The random forest ensemble training procedure was then subjected to stratified 15-fold cross-validation to obtain cancer type probabilities for the training samples. These probabilities were then used to train the ensemble of isotonic regressions for calibrating the random forest probabilities (**Figure 1b**).

### Performance of CUPLR

To assess the performance of CUPLR, we used the cancer type predictions based on the isotonic regression calibrated cross-validation probabilities, as well as the predictions upon applying CUPLR to the held out test set (**Figure 1b**). Both training (n=6139) and test sets (n=681) had the same cancer type proportions. Based on cross-validation predictions (**Figure 2**), CUPLR could classify all samples with an accuracy (i.e. fraction correctly classified) of 91%. Similar performance was observed for the test set across cancer types, with an accuracy of 89% across all samples (**Supplementary figure 5**). Certain cancer types show differences in test and CV accuracy which was due to low sample sizes (**Supplementary figure 6**).

**Figure 2:**
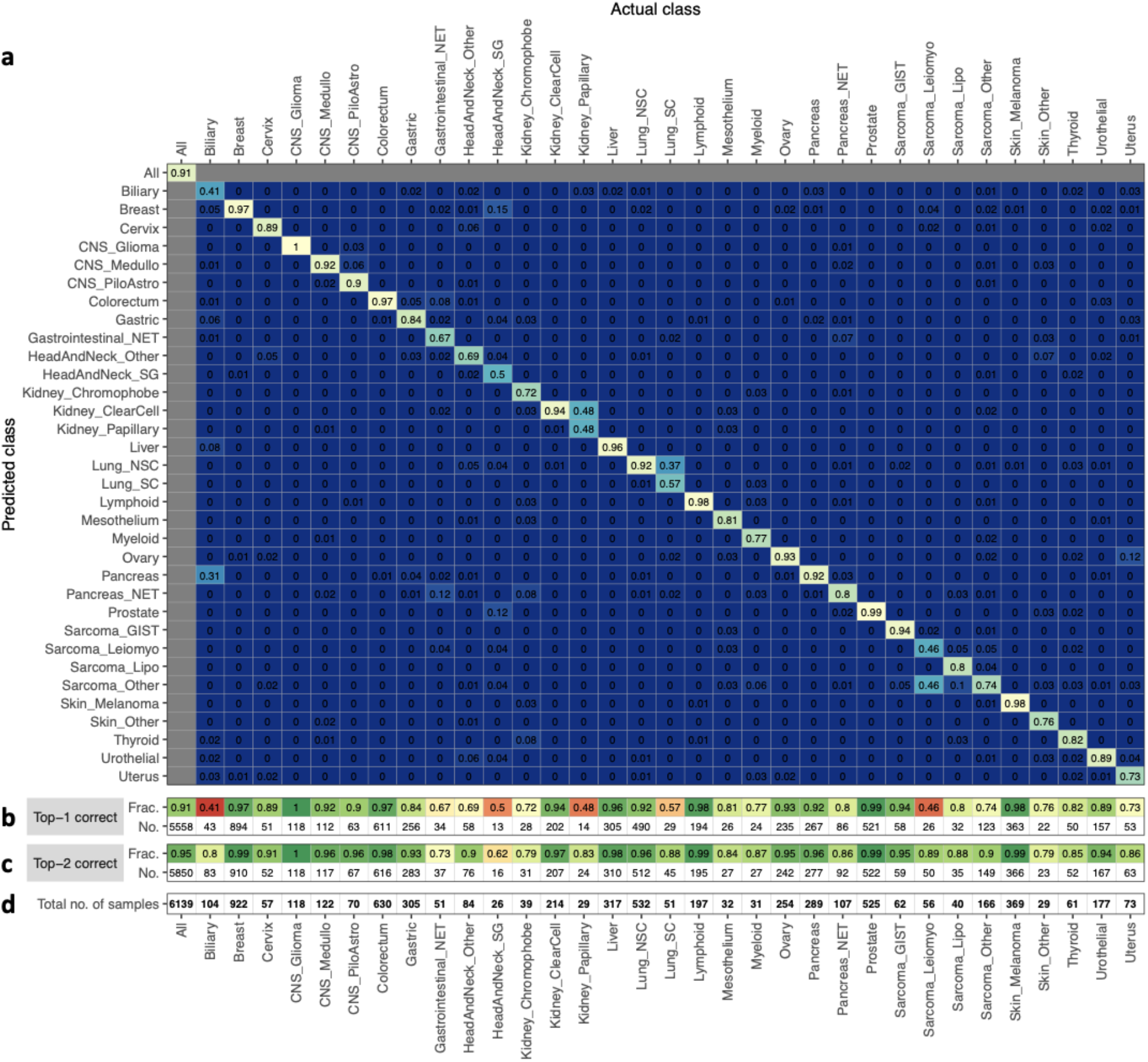
Accuracy of CUPLR based on cross-validation predictions. **(a)** A heatmap showing the accuracy of CUPLR where columns represent the fraction of samples in a cancer type cohort predicted as a particular cancer type. The diagonal represents the fraction of samples correctly predicted as a particular cancer type (i.e. accuracy). **(b)** The fraction and number of correctly classified samples, equivalent to the diagonal in **(a)**. **(c)** The fraction and number of correctly classified samples when considering the 2 highest probability cancer types. **(d)** The total number of samples for each cancer type. **Abbreviations**; *CNS*: central nervous system; *CNS_Medullo*: medulloblastoma; *CNS_PiloAstro*: pilocytic astrocytoma; *Gastrointestinal_NET*: gastrointestinal neuroendocrine tumor; *HeadAndNeck_SG*: salivary gland cancer; *HeadAndNeck_Other*: head and neck cancers other than salivary gland cancer; *Lung_NSC*: non-small cell lung cancer; *Lung_SC*: small cell lung cancer; *Pancreas_NET*: pancreatic neuroendocrine tumor; *Sarcoma_GIST*: gastrointestinal stromal tumor; *Sarcoma_Leiomyo*: leiomyosarcoma; *Sarcoma_Lipo*: liposarcoma; *Sarcoma_Other*: sarcomas other than leiomyosarcoma, liposarcoma or gastrointestinal stromal tumors; *Skin_Other*: non-melanoma skin cancer.

High misclassification rates for certain cancer types could likely be explained by shared cancer type characteristics (**Figure 2a**). This could be due to a common developmental origin, such as with Uterus being misclassified as Ovary (12%) due to both being gynecological cancers [22], and Biliary being misclassified as Pancreas (31%) and Liver (8%) due to being cancers of the foregut [23,24]. Cancer subtypes were also often misclassified as other subtypes, which was the case for Lung_SC (small cell lung cancer) towards Lung_NSC (non-small cell lung cancer) (37%); Kidney_Papillary towards Kidney_ClearCell (48%); and Sarcoma_Leiomyo (46%) and Sarcoma_Lipo (10%) towards Sarcoma_Other (sarcomas other than leiomyosarcoma, liposarcoma or gastrointestinal stromal tumors). Lastly, Gastrointestinal_NET (gastrointestinal neuroendocrine tumors) was occasionally misclassified as Pancreas_NET (pancreatic neuroendocrine tumors) (12%) which may, at least partially, reflect cancer type mis-annotation amongst these samples due to neuroendocrine tumors having similar morphological features and shared anatomical location [25].

Thus far, we have assessed performance based on whether the highest probability cancer type is the correct cancer type (i.e. top-1 accuracy; **Figure 2a,b**). However, if we consider whether the correct cancer type is amongst the top-2 highest probabilities (i.e. top-2 accuracy; **Figure 2c**), overall accuracy increases from 91% to 95%, with the greatest increases being for the cancer subtypes including Lung_SC (57% to 88%), Kidney_Papillary (48% to 83%), and Sarcoma_Leiomyo (46% to 89%). A large gain in accuracy was also observed for Biliary (41% to 80%) which was often misclassified as Pancreas. Similar increases in accuracy were seen based on predictions on the held out test set (**Supplementary figure 7**). The top-2 (and even top-3) probabilities of CUPLR are particularly useful for differential diagnosis purposes to narrow down potential TOOs when routine diagnostics are not fully conclusive.

### Added predictive value of SV related features

When examining the most important feature types from each random forest within CUPLR (**Figure 3a**), RMD profiles (‘rmd’) were consistently the most predictive of cancer type (in line with the findings from Jiao *et al.* 2020 [4]), as well mutational signatures (‘sigs’) including those with known cancer type associations such as SBS4 (associated with smoking [14]) in lung cancer (**Figure 3b**). As these mutation features are derived from genome-wide SBSs and indels, we assessed whether the presence of certain confounding factors that affect the SBS and indel genomic landscape (including DNA repair deficiencies [26,27], chemotherapy treatment [28,29], smoking history [30]) may lead to more incorrect predictions. However, these confounding factors showed minimal impact on classification performance (**Supplementary notes, Supplementary table 1, Supplementary figure 12**).

**Figure 3:**
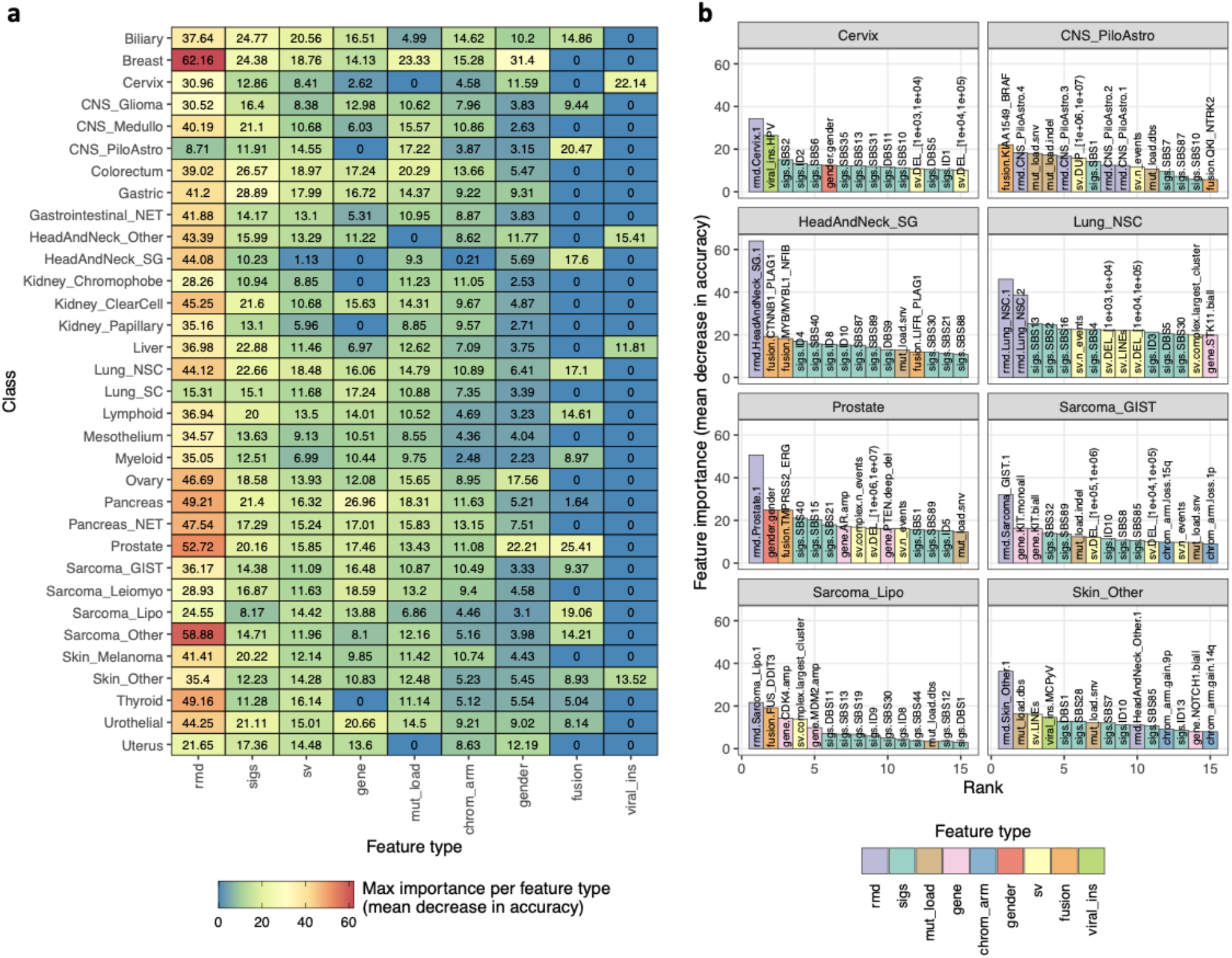
Features most predictive of cancer type. **(a)** Maximum feature importance per feature type for each cancer type random forest from CUPLR. **Feature type definitions**; rmd: regional mutation density profiles; sigs: mutational signatures; mut_load: total number of single base substitutions, double base substitutions or indels, gene: presence of gene gain or loss of function events; chrom_arm: chromosome arm copy number difference compared to overall genome ploidy; gender: sample gender as determined by sex chromosome copy number; sv: structural variants; fusion: presence of gene fusions; viral_ins: presence of viral sequence insertions. For a full description of each feature, see **Supplementary data 3**. **(b)** Importance of the top 15 features for selected cancer type random forests. Feature names are in the form {feature type}.{feature name}.

In addition to RMD profiles and mutational signatures, gender (as expected) was highly important for predicting cancers of the reproductive organs including breast, cervical, ovarian, prostate, and uterine cancer (**Figure 3b, Supplementary figure 8**). Notably, SV related features were important for classifying certain cancer types (**Figure 3b**). This included known cancer type specific gene fusions such as *TMPRSS2-ERG* in Prostate [31], *KIAA1549-BRAF* in CNS_PiloAstro (pilocytic astrocytomas) [32], *CTNNB1-PLAG1* and *MYBL1-NFIB* in HeadAndNeck_SG (salivary gland cancer) [33,34], and *FUS-DDIT3* in Sarcoma_Lipo (liposarcoma) [35]. Of note, while *EML4-ALK* fusions are known to be specific to non-small cell lung cancer [36], this fusion does not appear amongst the top 15 most important features in **Figure 3b** since it has low feature importance (**Supplementary data 4**) due to only occurring in a minor proportion of non-small cell lung cancer samples (**Supplementary figure 11**). We also find known viral DNA integrations as important features, including human papillomavirus (viral_ins.HPV) in Cervix [37] and Merkel cell polyomavirus (viral_ins.MCPyV) in Skin_Other (non-melanoma skin cancer) [38]. Lastly, the number of SVs in the largest complex SV cluster (sv.complex.largest_cluster) which we use as a proxy for the presence of chromothripsis was also predictive for liposarcomas, which are known to frequently harbor chromothripsis events. However, in contrast to the features mentioned above, the presence of chromothripsis alone is not sufficient for classifying a tumor as liposarcoma as chromothripsis is also highly prevalent in other tumor types [16].

To further assess the added value of SV related features, we excluded entire feature types from the training and examined the decrease in classifier performance (**Supplementary figure 9**). Indeed, removing gene fusion features (‘fusion’) resulted in a large drop in accuracy for HeadAndNeck_SG (salivary gland cancer; 43% to 23%) and Sarcoma_Lipo (liposarcoma; 80% to 63%), with removal of the viral integration features (‘viral_ins’) leading to decreased accuracy for Cervix (88% to 65%). Exclusion of simple and complex SV features (‘sv’) led to lower accuracy for Sarcoma_Lipo (80% to 73%) as well as Sarcoma_Leiomyo (leiomyosarcoma; 50% to 45%), likely due to the presence of chromothripsis (sv.complex.largest_cluster; **Figure 3b**) being important for the separation of these two cancer types, which are often misclassified as each other (**Figure 2a**). Similarly, removal of the simple and complex SV features (‘sv’) resulted in a large drop in accuracy for Lung_SC (small cell lung cancer; 53% to 41%) (**Supplementary figure 9**), likely because 1-10kb deletions and 10-100kb duplications distinguish Lung_NSC from Lung_SC (**Figure 3, Supplementary figure 8**).

When compared to existing WGS-based CUP classifiers (**Supplementary figure 10**), CUPLR also showed improved performance for cancer types where SV related features were important, including for CNS_PiloAstro, Lung_NSC, and Prostate. While Sarcoma_Lipo had no comparable cancer type in existing classifiers, prediction of all sarcoma subtypes by CUPLR (except for Sarcoma_Leiomyo) outperformed osteosarcoma prediction by PCAWG consortium classifier [4] (PCAWG-NN; Bone-Osteosarc) and sarcoma prediction from the Salvadores *et al.* [5] classifier (Salvadores-SVM; SARC). Overall, CUPLR performs similarly for the remaining cancer types (except for head and neck, myeloid and pancreatic neuroendocrine and thyroid cancers) when compared to existing classifiers, with CUPLR also being able to classify more cancer (sub)types.

In summary, the incorporation of all feature types resulted in the best performance, with SV related features being highly important for specific cancer types. To our knowledge, we are the first to show the added value of SV related features for cancer type classification.

### Graphical prediction report

Aside from cancer type probabilities, CUPLR also outputs explanations as to which features support these probabilities. These allow one to verify the predictions based on existing knowledge, and could be included in diagnostic reports to support decision making, e.g. in molecular tumor boards. The feature explanations are based on feature contribution calculations which enable feature importance determination at the sample level (rather than at the cohort level as in **Figure 3**). Specifically, feature contributions represent the total gain (or loss) in probability upon passing a feature through a random forest [39]. To ease the interpretation of CUPLR’s output, we have implemented a graphical report (**Figure 4**) which can be generated per patient that shows the cancer type probabilities (left panel), feature contributions for the top features for the top cancer types (middle panel). Also shown are the corresponding feature values in the patient in relation to the average feature value amongst patients in the training set (right panel).

**Figure 4:**
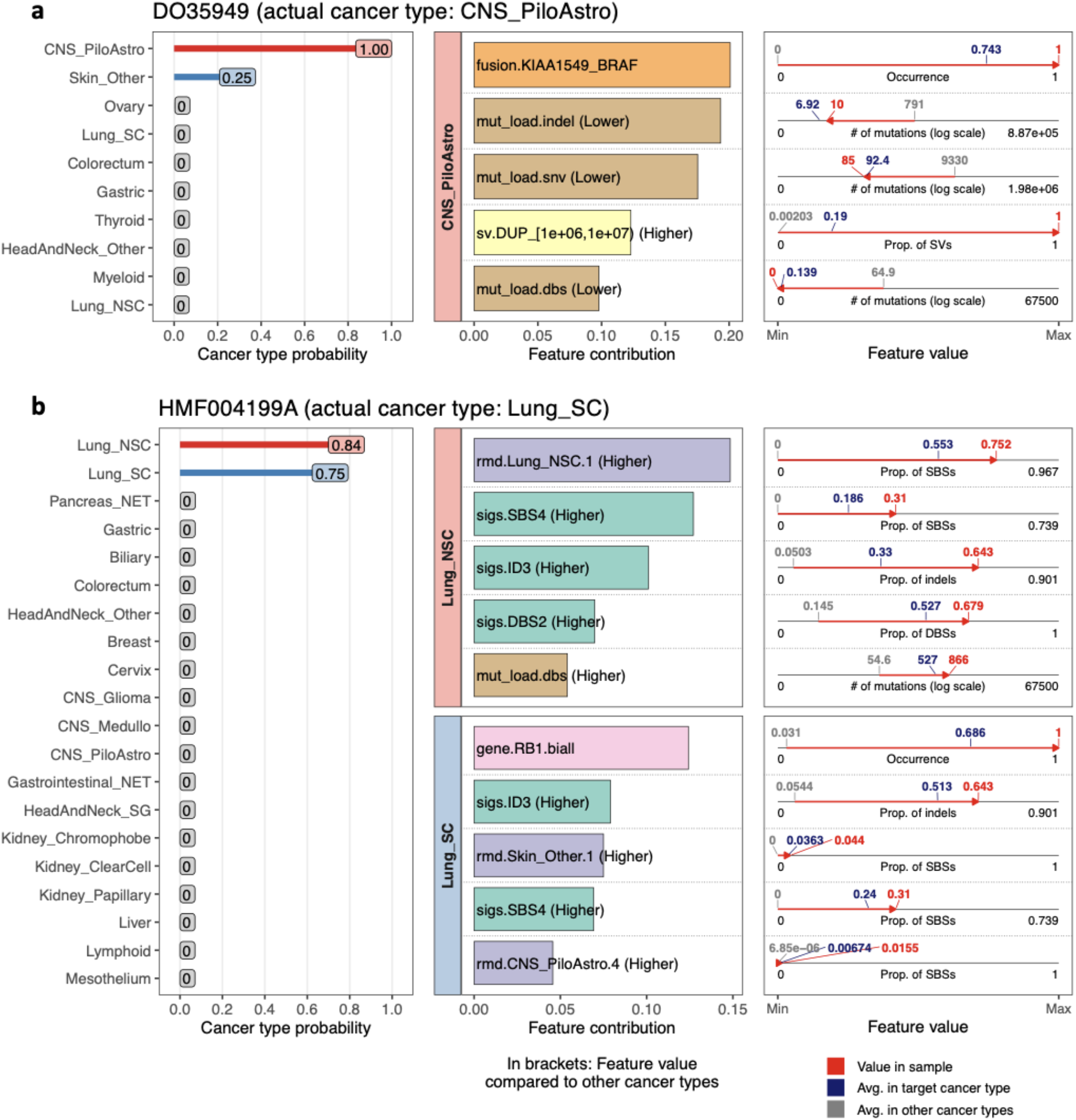
Graphical report for CUPLR predictions. Example reports are shown for two patients from the holdout set: **(a)** DO35949, annotated as having pilocytic astrocytoma (CNS_PiloAstro), and **(b)** HMF004199A, annotated as having small cell lung cancer (Lung_SC). The leftmost panels show the predicted cancer type probabilities. In the middle panels, contributions of the top features are shown for the top predicted cancer types. When there is uncertainty in the cancer type probabilities such as in **(b)**, more feature contribution panel rows are shown. In the rightmost panel, the feature values in the patient are plotted in relation to the average feature value amongst patients in the training set with and without the target cancer type. For non-Boolean features (i.e. not those where the values are present or absent), the red arrows indicate whether the feature value in the patient is higher or lower than in patients without the target cancer type. For a full description of each feature, see **Supplementary data 3**.

We will use patient DO35949 as an example demonstration of the graphical report (**Figure 4a**). Since the CNS_PiloAstro probability (1.00) was much higher than the probability of other cancer types, only one cancer type (i.e. panel row) is shown, with up to 3 panel rows being shown when more than one cancer type probability is high (such as for patient HMF004199A; **Figure 4b**). The top feature for DO35949 was the presence of a KIAA1549-BRAF fusion (middle panel) which is on average found in 74% of CNS_PiloAstro patients (blue label), but 0% in all other patients (grey label). Since this feature is of Boolean type, a feature value of 1 (red label) indicates the presence of the *KIAA1549-BRAF* fusion in DO35949 (whereas a feature value of 0 would indicate absence). On the other hand, the mutational load of SNVs, DBSs and indels (mut_load.snv, mut_load.dbs, mut_load.indel) is on average lower in DO35949 (red label) and CNS_PiloAstro patients (blue label) compared to all other patients (grey label). Conversely, the proportion of 1-10Mb duplications (sv.DUP_[1e+06,1e+07]), is on average higher in DO35949 (red label) and CNS_PiloAstro patients (blue label) compared to all other patients (grey label). For all features, a red arrow (going right or left)highlights whether the feature value in DO35949 is higher or lower (respectively) than that in all other patients. Additionally, this relationship is also indicated in text in the feature contribution panel (middle panel) for non-Boolean features.

Patient HMF004199A (**Figure 4b**) is an example of a situation where more than one cancer type probability is high, with the probability of Lung_NSC (non-small cell lung cancer) being 0.84 and Lung_SC (small cell lung cancer) being 0.75 (**Figure 4b**). Here, two feature panels are shown for both of these cancer types to aid with resolving this uncertainty. Since *RB1* loss is specific to small cell lung cancer and rarely occurs in non-small cell lung cancer [40,41], it is more likely that the correct predicted cancer type is Lung_SC and not Lung_NSC. Indeed, this patient was diagnosed with small cell lung cancer.

This graphical report presents the detailed machine learning-based classification output of CUPLR in a human readable format. We acknowledge that the output is highly detailed, which is inevitable due to the large amounts of data used by the algorithm. However, as these details may not be necessary in all circumstances, we have implemented the option in the software to hide the feature contribution and/or feature value panels in the final graphical output.

### Feature contributions aid in clarifying the primary tumor location of CUPs

To showcase how CUPLR could be used in a real life clinical setting, we applied CUPLR to 149 tumors for which primary tumor location was unknown from the Hartwig dataset, and examined the top predicted cancer type together with the top contributing features for each sample (**Supplementary data 2**). Of the 75 samples with a probability ≥0.8 for the top cancer type, the cancer type for 29 of these samples could be confidently determined purely based on these samples exhibiting features with well-known cancer type associations. This included 22 samples with signatures of smoking (SBS4 and ID13 [14]) and predicted as Lung_NSC, 1 sample with signatures of ultraviolet radiation (SBS7, DBS1 [14]) exposure and predicted as Skin_Melanoma, 5 samples with *APC* mutations and predicted as Colorectum [42], and 1 sample with a *TMPRSS2-ERG* fusion [31] and predicted as Prostate.

An additional 15 of the 75 samples could be confidently annotated based on having known cancer type specific features in conjunction with having a high contribution of the RMD profile of the predicted cancer type. For example, *KRAS* mutations most commonly occur in pancreatic cancer but are also frequent in other cancer types such as colorectal or lung cancer (**Supplementary figure 11**) [43]. For 3 samples with a *KRAS* mutation and predicted as Pancreas, the top contributing feature was rmd.Pancreas.1 (RMD profile of pancreatic cancer) thereby giving us confidence that the Pancreas classification was correct. Likewise, in conjunction with the respective RMD profile, 10 samples could be clarified as Gastric based on the presence of the signature of ROS damage (SBS17) [14], LINE insertions [44], and/or fragile site deletions [45]; 1 sample as HeadAndNeck_Other based on having HPV integration [46]; and 1 sample as Uterus based on having a *PIK3R1* mutation [47] (**Supplementary data 2**). For the remaining 31 of the 75 samples, no known cancer type specific features could be identified, although the RMD profiles were the primary contributors of the top cancer type probability for these samples. Most of these samples were predicted as cancer types with highly specific RMD profiles respective to each cancer type, including Breast, Colorectum, Liver, Kidney_ClearCell, Ovary, and Urothelial (**Supplementary data 2**, **Supplementary figure 11**), and classification of cancer types achieved ≥0.9 accuracy (**Figure 2a**). The majority of these samples are thus likely correctly annotated, although additional evidence would be required for validation and final diagnosis. This could for example be based on histopathological examinations, but these were unfortunately not available for the retrospectively analyzed samples that were included here. However, such information would be readily available in routine diagnostics.

## Discussion

Here, we have developed a tissue of origin classifier (CUPLR) for the analysis and diagnostics of CUPs using a large uniformly processed WGS dataset of both primary and metastatic tumors. Our classifier is to our knowledge the first to incorporate genomic features derived from SVs, as well as provide human interpretable explanations alongside each prediction which allows for manual resolving of CUPs, especially in cases of lower scoring samples or for samples for which multiple tumor types are suggested by CUPLR.

Current state-of-the-art WGS-based classifiers, including those by Jiao *et al.* [4] and Salvadores *et al.* [5], achieve high accuracy (≥80%) for over three quarters of the cancer types they predict by primarily employing RMD and mutational signatures as features, which are derived from simple mutations. CUPLR builds upon these classifiers, with the inclusion of SV and driver gene related features improving performance for certain cancer types, such as for pilocytic astrocytoma and prostate cancer. Of note, genomic-based TOO classifiers published thus far have only used data from primary tumors [4,5,12,13]. However, since CUPs are by definition metastatic tumors, the inclusion of 4401 metastatic tumors with known tissue of origin for training CUPLR may make it a more suitable tool for the purpose of clarifying CUPs. We do acknowledge however that the metastatic tumor data used here may harbor treatment effects which are absent in treatment-naive CUPs, such as driver mutations in *AR* [48] or *ESR1* [49] conferring resistance to therapy or presence of characteristic mutations induced by for example 5-fluorouracil [28] or radiotherapy [29,50]. Identification and removal of such treatment associated features could potentially improve TOO classification. Additionally, CUPLR is able to distinguish the largest number of cancer (sub)types (33 cancer types) compared to existing WGS-based classifiers, with the Jiao [4] and Salvadores [5] classifiers discriminating 24 and 18 cancer types respectively, and also more than a recently published whole histology slide image based classifier which discriminates 18 cancer types [51]. We do acknowledge that discriminating even more cancer types is warranted, but this is currently limited by the amount of available training samples that were sequenced and uniformly bioinformatically analyzed. For example, thymic cancer or lung neuroendocrine tumors had too few (<20) samples to be included as separate classes for training (**Supplementary data 1**). Furthermore, certain cancer types could be divided into their subtypes, such as Myeloid (currently only with 31 samples; **Figure 2a**) into acute myeloid leukemia and multiple myeloma [52], and sarcomas in a broader range of subtypes [53]. The availability of more WGS data for less frequent, but also (ultra-)rare cancers would allow for training of an updated CUPLR model that classifies additional cancer types and subtypes.

While CUPLR achieved overall excellent classification accuracy (91%), which is similar to or better than other WGS-based classifiers (**Supplementary figure 10**), it should be noted that this is driven by several large sub-cohorts of common cancer types (e.g. breast and colorectal cancer) and that performance for certain cancer types is still suboptimal. For cancer subtypes that CUPLR has difficulty discriminating, such as small cell versus non-small cell lung cancer and papillary versus clear cell renal carcinoma, additional information from histopathological staining and immunohistochemistry could be used to clarify these cases. Here, the application of artificial intelligence-based histology image analysis [51] could further improve the accuracy and reliability of resolving CUPs. Clinical metadata, such as biopsy location and metastasis grade [54], when used together with CUPLR can also aid with clarifying primary tumor location. For instance, there may be uncertainty as to whether a tumor with human papillomavirus DNA integration was a head and neck cancer or cervical cancer based on CUPLR predictions (e.g. DO15430, **Supplementary data 2**), since human papillomavirus DNA integration is characteristic of these two cancer types [37]. However, in clinical practice, it will always be known if the tumor biopsy was taken from the upper body and if the lesions were local, to be able to conclude that this tumor can only be of head and neck origin.

Given that RMD [4,5], mutational signatures [55], and SVs [20,56] are still active areas of research, we expect improvements to these features could also lead to better TOO classification. Here, we have demonstrated that extraction of cancer type specific RMD profiles is possible from raw mutation counts, similarly as was done for mutational signatures [14]. However, CUPLR does not heavily rely on RMD profiles for classification of certain cancer types such as liposarcoma and non-melanoma skin cancer (**Figure 3b**), potentially because more training samples are required to extract more stable and informative RMD profiles which could improve classification for these cancer types. Improvements to TOO classification may also come from extraction of more comprehensive mutational signatures, for example by incorporating information of mutation timing or genome localization [57,58]. Development of more sophisticated signature extraction methods may also allow for quantification of low signal tissue type specific signatures, such as SBS88 (associated with colibactin induced DNA damage in the colon) which has only been extracted from colon healthy tissue [59,60] but not cancer tissue likely because other mutational processes in cancer tissue mask the presence of this signature (**Supplementary figure 8**, **Supplementary figure 11**). Lastly, while CUPLR uses a wide range of features derived from SVs (including gene fusions, viral DNA integrations, LINE insertions, structural deletion and duplication size, and chromothripsis), there is still an opportunity to explore other SV related features to improve TOO classification, such as SV signatures [56].

Aside from WGS data, gene expression [6,7,61] and methylation [8] data have also been used to develop TOO classifiers, and inclusion of such data could further improve cancer type classification, although there may be strong redundancy with RMD information derived from WGS. As application of gene expression and methylation profiling for cancer diagnostics is still in its infancy [9,10] and the inclusion of additional procedures would likely increase costs and time, its potential added value should thus be well balanced against these extra burdens. Furthermore, reliable gene expression profiling depends on the integrity and quality of RNA [62], which remains a challenge for routine clinical samples. Thus, to facilitate clinical implementation of CUPLR in diagnostic centers that have started implementing WGS in routine cancer diagnostics, the model was trained solely on WGS data despite RNA-seq data being available for subsets of samples of both the Hartwig [63] and PCAWG datasets [43].

Given the gradually increasing adoption of WGS in routine diagnostics [11], CUPLR serves as a viable complementary tool to standard procedures for determining TOO (e.g. histopathological staining and immunohistochemistry). CUPLR can be run from the output of open source tools and is freely available as an R package at https://github.com/UMCUGenetics/CUPLR. The trained model as well as the code for generating the input features is provided to enable prediction on new samples and for facilitating diagnostic use.

## Methods

### Datasets

We have used whole genome sequencing data from cancer patients from two cohorts: the Hartwig Medical Foundation (Hartwig) cohort and the Pan-Cancer Analysis of Whole Genomes (PCAWG) cohort.

The Hartwig cohort contained patient data for which re-use for cancer research was consented by the patients as part of clinical studies (NCT01855477, NCT02925234) that were approved by the medical ethical committees (METC) of the University Medical Center Utrecht and the Netherlands Cancer Institute, and was provided under data transfer agreement (DR-104) by Hartwig Medical Foundation. The Hartwig cohort included 4776 metastatic tumor samples from 4635 patients. For patients with multiple biopsies that were taken at different timepoints, patient IDs were suffixed by ‘A’ for the first biopsy, ‘B’ for the second biopsy, etc (e.g. HMF001423A, HMF001423B).

The PCAWG cohort consisted of 2650 patient tumors, and access for raw sequencing data for these tumors was granted via the Data Access Compliance Office (DACO) Application Number DACO-1050905 on 6 October 2017 and via https://console.cancercollaboratory.org download portal on 4 December 2017.

Sample selection for developing CUPLR were based on several criteria. First, for patients with multiple biopsies, the biopsy with the highest tumor purity was selected. Second, samples with <0.2 tumor purity were removed as somatic variant calling was not reliable for these samples. Third, PCAWG samples that have been grey- or blacklisted by the PCAWG consortium were removed (https://dcc.icgc.org/releases/PCAWG/donors_and_biospecimens). Fourth, samples with low mutational load were removed (≤35 single base substitutions), except for medulloblastoma and pilocytic astrocytoma samples. Lastly, cancer type cohorts with <20 samples were removed.

### Variant calling

Variant calling for both the Hartwig and PCAWG cohorts was performed using the Hartwig Medical Foundation Platinum pipeline (v5) (https://github.com/hartwigmedical/pipeline5). Briefly, reads were mapped to GRCh37 using BWA (v0.7.17). GATK (v3.8.0) Haplotype Caller was used for calling germline variants in the reference sample. SAGE (v2.2) was used to call somatic single and multi nucleotide variants as well as indels. GRIDSS (v2.9.3) was used to call simple and complex structural variants. PURPLE (v2.53) combines B-allele frequency (BAF) from AMBER (v3.3), read depth ratios from COBALT (v1.7), and structural variants from GRIDSS to estimate copy number profiles, variant allele frequency (VAF) and variant clonality. PURPLE also determines sample gender based on sex chromosome ploidy. LINX (v1.14) interprets structural variants (to identify simple and complex structural events) from PURPLE, and also detects gene fusions, viral DNA integrations, and homozygously disrupted genes.

### Extraction of features

#### Regional mutational density

RMD was defined as the number of somatic SBSs in each 1Mb bin across the genome (n=3071), normalized by the total number of SBSs in the sample. Extraction of RMD profiles from these RMD bins was performed within the CUPLR training procedure (**Supplementary figure 1a**) using non-negative matrix factorization (NMF). For details, see **Methods: CUPLR training procedure**.

#### Mutational signatures

The number of somatic mutations falling into the 96 SBS, 78 DBS and, 83 indel contexts (as described in COSMIC: https://cancer.sanger.ac.uk/signatures/) was determined using the R package mutSigExtractor (https://github.com/UMCUGenetics/mutSigExtractor, v1.23). To obtain the mutational signature contributions for each sample, the mutation context counts were fitted to the COSMIC catalog of mutational signatures using the *nnlm()* function from the NNLM R package.

The contributions of the child signatures SBS7a, SBS7b, SBS7c and SBS7d were summed to yield the parent signature SBS7. Similarly SBS10a-d and SBS17a-b were merged to yield SBS10 and SBS17. Lastly, the SBS, DBS and indel signature contributions were normalized by the total number of SBSs, DBSs and indels respectively.

#### Chromosome arm and overall genome ploidy

Chromosome arm and genome ploidies were determined in a similar method as described by Taylor *et al.* 2018 [18].

Somatic copy number (CN) segments (called by PURPLE) were split by their respective chromosome arms. Only chr1-22 and chrX were included. All chromosomes have *p* and *q* arms, except for chr13, chr14, chr15, chr21, and chr22 which are considered to only have the *q* arm. For each chromosome arm, the CN values of each segment were converted to integer values. The arm coverage of each CN integer value was then determined (e.g. 70% of the arm has a CN of 2, 20% a CN of 1 and 10% a CN of 3). The CN with the highest coverage was assigned as the preliminary arm CN. The most common CN across all arms was assigned as the genome CN.

Two filtering steps were then performed to obtain the final arm CN values. For each arm, if the CN with the highest coverage has <50% coverage, and if any of the CN values equal the genome CN, then assign that CN as the final CN of the arm. Else, assign the genome CN as the final CN of the arm.

To determine the CN gains and losses of each arm, the difference between the arm CN and genome CN was calculated. Arms with CN differences >1 were considered to have a gain, and <1 a loss.

#### Features extracted from LINX output

LINX combines simple mutations (point mutations and indels), structural variants, and copy number variants to resolve simple and complex structural rearrangements, and subsequently identify gene driver events, gene fusions and detect viral DNA integrations.

The presence of 4 types of gene driver events were determined from the output of LINX: (i) amplifications, (ii) deep deletions, (ii) biallelic loss, and (iv) monoallelic hits (pathogenic mutation in one allele). Amplified genes were marked as *likelihoodMethod==’AMP’* by LINX. Genes with deep deletions were marked as *likelihoodMethod==’DEL’*. Genes with biallelic loss were marked as *biallelic==TRUE*. Genes with a monoallelic pathogenic mutation were marked as *biallelic==FALSE* and *driver==”MUTATION”*, and we selected only those with *driverLikelihood>=0.9* (referring to the likelihood of the mutation being impactful as determined by the dndscv R package [64]). LINX determines the presence of driver events for 462 genes. Thus, there are 462 genes x 4 driver types = 1848 gene driver features in total. We then performed preliminary feature selection to reduce the computational resources required for training CUPLR. Here, one-sided Fisher’s exact tests were performed and Cramer’s V values were calculated. Only genes where at least one driver type had a p-value <0.01 and Cramer’s V ≥0.1 were kept (205 genes x 4 driver type = 820 gene driver features).

Gene fusions belonged to 3 categories: (i) well known fusion pairs (e.g. *TMPRSS2-ERG*), (ii) immunoglobulin heavy chain (IGH) locus fusions, and (iii) fusions with promiscuous gene partners (e.g. *BCR*). *IGH* fusions were grouped into a single feature as these are characteristic events in lymphoid cancers [65]. Fusions with one promiscuous gene partner were grouped by gene (e.g. *RUNX1_ETS2* and *RUNX1_RCAN1* would both fall under the *RUNX1_\** feature). Fusions with two promiscuous gene partners were split into two separate features (e.g. *SLC45A3_MYC* would become the features *SLC45A3_\** and *\*_MYC*). Only fusions that were marked as *reported==TRUE* by LINX (i.e. reported in literature) were selected. We then performed preliminary feature selection due to the large number of possible fusions present in our dataset (n=512). Here, one-sided Fisher’s exact tests were performed and Cramer’s V values were calculated. Only fusions with p-value <0.01 and Cramer’s V >=0.1 were kept (45 fusions).

For the viral DNA integrations present in our dataset, we merged virus strains into 9 virus categories: adeno-associated virus (AAV), Epstein-Barr virus (EBV), hepatitis B virus (HBV), hepatitis C virus (HCV), human immunodeficiency virus (HIV), human papillomavirus (HPV), herpes simplex virus (HSV), human T-lymphotropic virus (HTLV), and Merkel cell polyomavirus (MCPyV). For example, *human papillomavirus type 16* and *human papillomavirus type 18* would be both grouped as *human papillomavirus*.

LINX chains individual SVs into SV clusters and classifies these clusters into various event types. Clusters can have one SV (for simple events such as deletions and duplications), or multiple SVs. We defined SV load as the total number of SV clusters. We quantified deletions and duplications (*ResolvedType is ‘DEL’ or ‘DUP’*) stratified by length (1–10 kb, 10–100 kb, 100kb–1Mb, 1–10Mb, >10 Mb), as well as long interspersed nuclear element (LINE) insertions (*ResolvedType==‘LINE’*), and normalized these counts by SV load. We defined the number of double minutes as the number of clusters where *InDoubleMinute==TRUE*. We defined the number of foldback events as the number of clusters where *InfoStart==‘FOLDBACK’ or InfoEnd==‘FOLDBACK’ or ResolvedType==‘RESOLVED_FOLDBACK’*. For clusters where *ResolvedType==‘COMPLEX’*, we defined the complex SV load as the total number of these clusters, and the largest complex SV cluster as the cluster with most SVs.

### CUPLR training procedure

#### Extraction of regional mutational density profiles

To extract the cancer specific RMD profiles from the 3071 RMD bins, a multistep procedure involving non-negative matrix factorization (NMF) (**Supplementary figure 2**) was performed prior to classifier training (**Supplementary figure 1ai**).

All NMF runs described below are performed with the *nnmf()* function from the NNLM R package (v0.4.4) with the parameters *loss=’mkl’* and *max.iter=2000*.

For each cancer type cohort, an NMF rank search was done to determine the optimum rank (i.e. number of RMD profiles) (**Supplementary figure 2a**). For ranks 1 to 10, NMF was performed 50 times on a random subset of 100 samples from the cohort (or if the cohort contained less than 100 samples, all samples from that cohort were used) with 10% of the values randomly removed. The missing values were then imputed and the mean squared error (MSE) of these imputed values was calculated. This method of calculating MSE is described by the authors of the NNLM R package [66]. The median of the MSE across the 50 NMF iterations was then calculated. The rank search thus results in 10 MSE values across the 10 ranks searched. The optimum rank was the rank before the increase in log10(MSE) was >0.2%, and NMF was then performed using the optimum rank and without removing random values to produce the RMD profiles for the cancer type cohort (**Supplementary figure 2b**).

The above procedure thus yields a different set of RMD profiles for each cancer type cohort. However, some RMD profiles across related cancer types (e.g. pancreas and biliary cancer) may actually be equivalent RMD profiles. Hierarchical clustering (using Pearson correlation as a distance measure) was thus performed to group similar RMD profiles across all cancer type cohorts. The resulting dendrogram was cut at a height of 0.1 (using the R function *cutree()*), whereby RMD profiles under this height were grouped and considered the same profile. From each of the groups, one profile was greedily selected to yield the final set of RMD profiles (**Supplementary figure 2c**).

To obtain the RMD profile contributions for each sample, the RMD bins were fitted to the RMD profiles using the *nnlm()* function from the NNLM R package.

#### Random forest ensemble training

The main component of CUPLR comprises an ensemble of binary random forests that each discriminate one cancer type (**Supplementary figure 1aii**). The below text describes the training procedure for each cancer type random forest.

First, univariate feature selection was performed to remove irrelevant features (**Supplementary figure 1aiii**). Pairwise testing was done to compare feature values from samples of the target cancer type (case group) versus the remaining samples (control group). For numeric features, p-values were determined using Wilcoxon rank sum tests, and effect sizes were calculated using Cliff’s delta. For boolean features, p-values were determined using Fisher’s exact tests, and effect sizes were calculated using Cramer’s V. Depending on the feature, alternative hypotheses for the Wilcoxon rank sum tests and Fisher’s exact tests were one or two sided. See **Supplementary data 3** for details on which features are numeric or boolean, as well as which alternative hypothesis was used. Features were kept which had p<0.01 and effect size ≥0.1. The number of features kept was capped to 100 features.

Second, random oversampling was performed for the case group which always contained fewer samples than the control group, which were randomly undersampled (**Supplementary figure 1aiv**). A grid search was performed to determine the optimal pair of 4 oversampling and 4 undersampling ratios. These ratios were automatically determined as follows: i) calculate the geometric mean between the case and control group sample sizes; ii) the 4 resampling ratio values are logarithmically spaced between the geometric mean and the case group sample size or the control group sample size. For each over-/undersampling ratio pair, stratified 10-fold cross-validation (CV) was performed, after which the area under the precision recall curve (AUPRC) was calculated. The pair with the highest AUPRC was chosen and the resampling was applied.

Lastly, a random forest was trained (**Supplementary figure 1av**) using the *randomForest* R package (v4.6-14) with default settings. A filter is applied to the probabilities produced by the random forest based on sample gender, where breast, ovary and cervix probabilities are set to 0 for male samples, and prostate probabilities are set to 0 for female samples. Local increments were calculated for the random forest using the *rfFC* R package (v1.0) to enable downstream calculation of feature contributions [39].

#### Isotonic regression training

The entire random forest ensemble training procedure was then subjected to stratified 15-fold cross-validation which allows every sample to be excluded from the training set in order to obtain cancer type probabilities for the training samples (Supplementary figure 1b). These cross-validation probabilities were then used to train an ensemble of isotonic regressions using the *isoreg* R function (one per cancer type random forest) to calibrate the probabilities produced by the random forest ensemble (Supplementary figure 1c, Supplementary figure 3).

Random forests tend to be overconfident at probabilities towards 0 and underconfident at probabilities towards 1 [21], and this bias varies between random forests (**Supplementary figure 4**). In other words, a probability of e.g. 0.8 from one random forest does not correspond to a probability of 0.8 from another random forest. Probability calibration greatly reduced this bias ensuring that predictions across the random forests are comparable (**Supplementary figure 4**).

#### Performance evaluation

To assess the performance of CUPLR, we used the cancer type predictions based on the isotonic regression calibrated cross-validation probabilities, as well as by applying the final CUPLR model to a validation set whereby 10% of samples were held out from the full training set. Accuracy was calculated for each cancer type (as well as across all cancer types), which was the fraction of samples where the top predicted cancer type was the actual cancer type for the respective sample. Top-N accuracy was also calculated, which was the fraction of samples where the actual cancer type was amongst the top N predicted cancer types for the respective sample. ‘Accuracy’ is equivalent to ‘top-1 accuracy’.

## Data availability

Metastatic WGS data and corresponding metadata have been obtained from the Hartwig Medical Foundation and provided under data request numbers DR-104. Both WGS data and metadata is freely available for academic use from the Hartwig Medical Foundation through standardized procedures and request forms can be found at https://www.hartwigmedicalfoundation.nl. For access to identifying data (e.g. germline or raw read data) for the Pan-Cancer Analysis of Whole Genomes (PCAWG) cohort, researchers will need to request access via the ICGC Data Access Compliance Office (DACO; https://daco.icgc.org/). The extracted features for each sample and used to develop CUPLR is available at https://doi.org/10.5281/zenodo.5547228. All other data are available within the article, Supplementary Information or available from the authors upon request.

## Code availability

The Hartwig Medical Foundation Platinum (https://github.com/hartwigmedical/pipeline5) is used for germline and somatic variant calling, as well as post-processing procedures such as identification of simple and complex structural rearrangements, annotation of driver gene mutation events, and detection of gene fusions and viral DNA integrations. CUPLR can be run from the output of this pipeline, and is available as an R package at https://github.com/UMCUGenetics/CUPLR. The code used for data processing and generating the figures is also available in this repository.

## Acknowledgements

This research was supported by an unrestricted grant of Stichting Hanarth Fonds, The Netherlands. This publication and the underlying study have been made possible partly on the basis of the data that Hartwig Medical Foundation and the Center of Personalised Cancer Treatment (CPCT) have made available to the study.

## Author contributions

L.N. performed analyses, wrote/edited the paper. A.V.H. conceived the study, performed analyses, wrote/edited the paper. E.C. edited the paper and provided discussion. E.C. and A.V.H. supervised the study. All authors proofread, made comments, and approved the paper.

## Supplementary data

**Supplementary data 1**: Metadata for tumors in the Hartwig and PCAWG datasets, including cancer type label, HRD and MSI status, and inclusion/exclusion in the training and holdout sets.

**Supplementary data 2**: Output from CUPLR based on cross-validation predictions on the training set, and predictions on the holdout set and on cancer of unknown primary samples.

**Supplementary data 3**: Description of each feature used by CUPLR.

**Supplementary data 4**: Feature importance for each random forest within CUPLR.

## Supplementary information

### Supplementary figures

**Supplementary figure 1:**
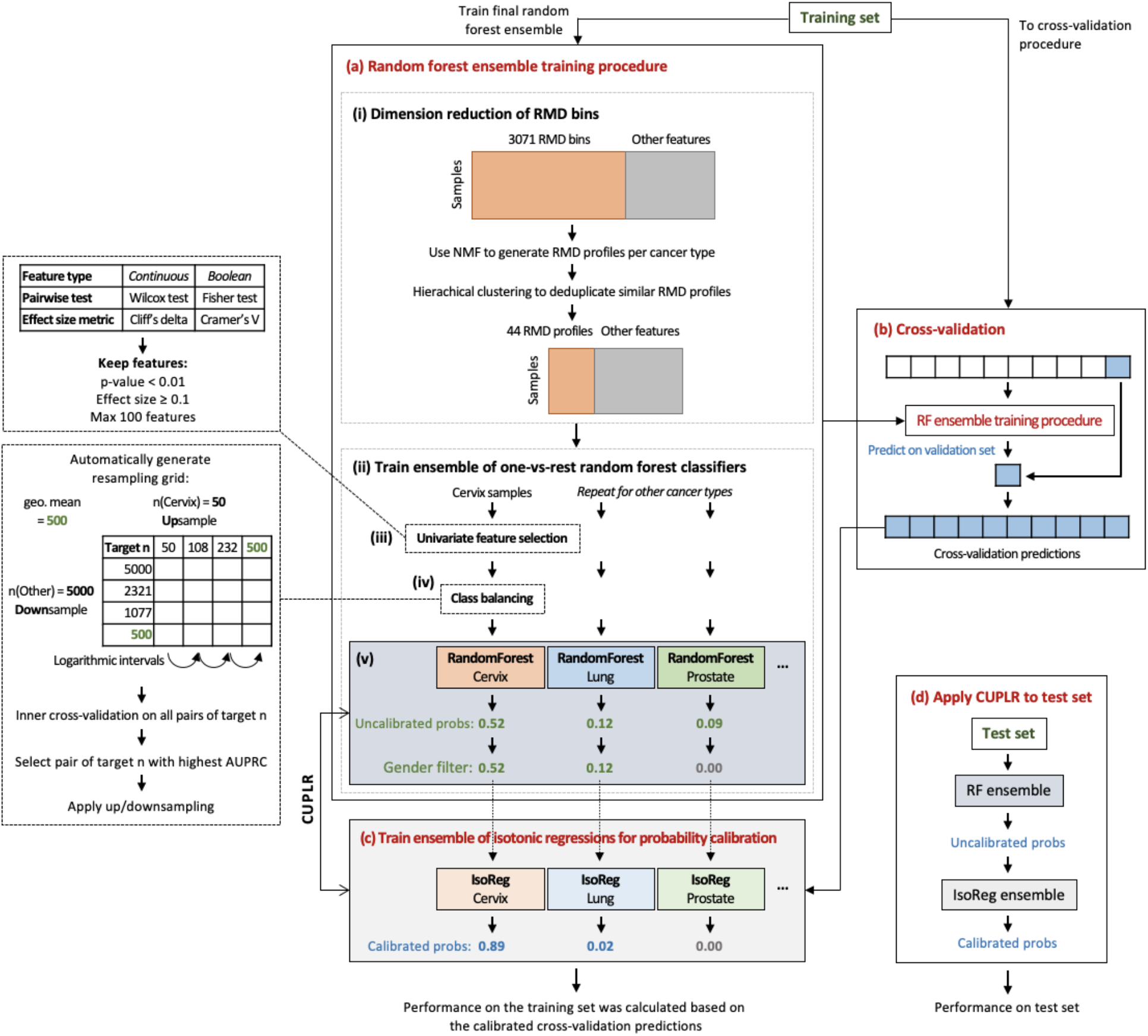
CUPLR training procedure. **(a)** An ensemble of binary random forest classifiers was trained each to discriminate one cancer type versus other cancer types. **(i)** Dimension reduction via non-negative matrix factorization (NMF) was performed on the 3071 RMD bins independently for each cancer type to ultimately produce 44 cancer type specific RMD profiles (see **Supplementary figure 2** for a detailed schematic), prior to random forest training as shown in **(ii)**. **(iii)** Univariate feature selection was performed to remove irrelevant features. (**iv**) Class resampling was performed to alleviate imbalances in the number of samples for each cancer type. (**v**) The random forests are trained. Breast, ovary and cervix probabilities from the random forest are set to 0 for male samples, and prostate probabilities are set to 0 for female samples. **(b)** The whole training procedure in **(a)** was subjected to 15-fold cross-validation to obtain cancer type probabilities for the training samples. **(c)** These probabilities were then used to train the isotonic regressions for probability calibration. The calibrated probabilities were also used to calculate cross-validation performance. **(d)** Performance was also determined by applying CUPLR to a held out test set.

**Supplementary figure 2:**
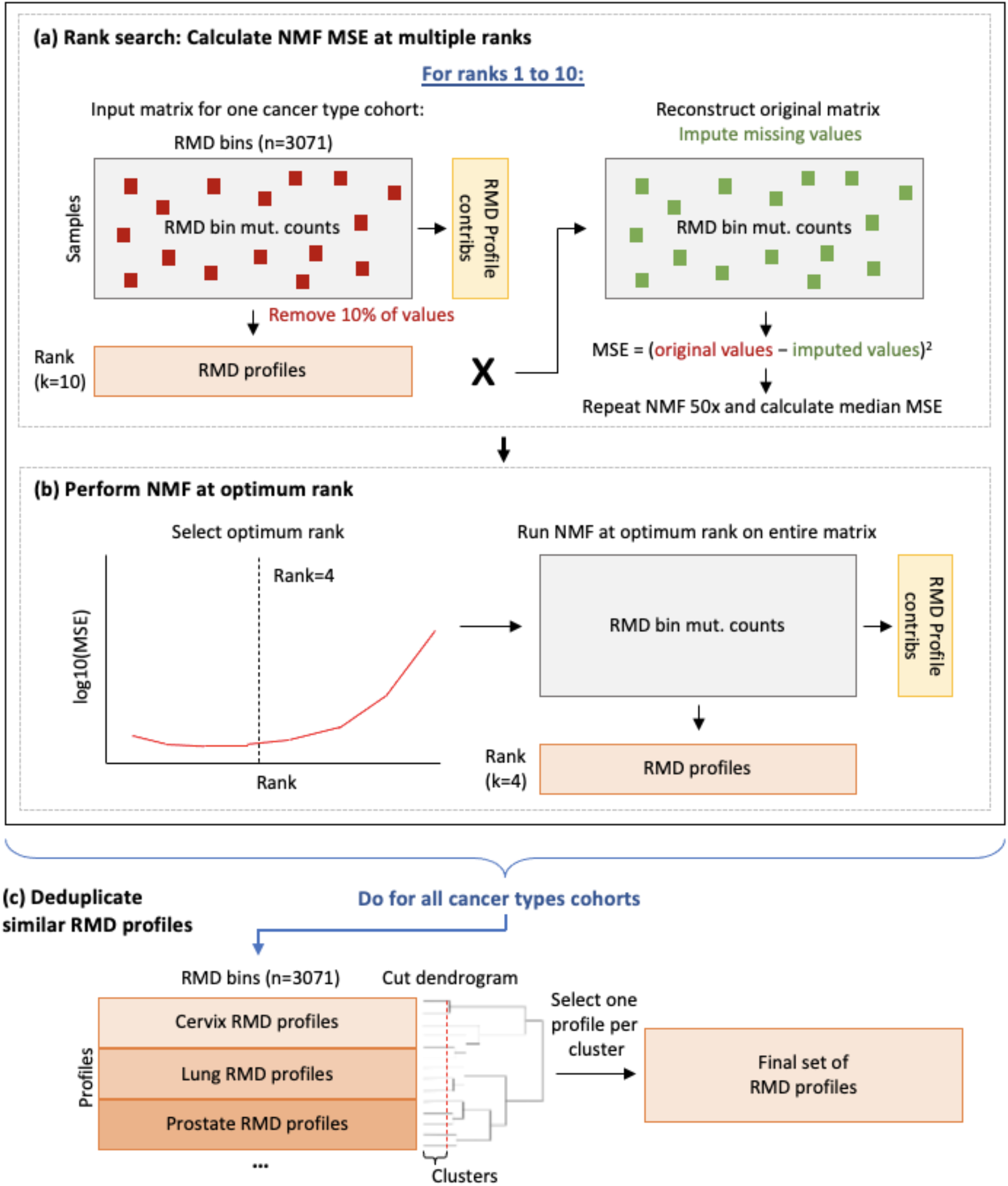
Extraction of regional mutational density (RMD) profiles. **(a)** Non-negative matrix factorization (NMF) was performed for several ranks to determine the optimum rank. For each rank, 10% of values were removed from the input matrix, NMF was performed, the original matrix was reconstructed, the missing values were imputed, and mean squared error between the original missing values and the imputed values. This was repeated 50 times and median MSE was calculated. **(b)** The optimum rank was the one at which the log10(MSE) value began to increase rapidly. NMF was performed on the entire input matrix (i.e. without removing values) to yield the RMD profiles for one cancer type cohort. **(c)** The procedures in (a) and (b) were performed for all cancer types to yield RMD profiles for all cancer types. Hierarchical clustering was performed to group profiles that were similar. One profile was selected per cluster to yield the final set of RMD profiles.

**Supplementary figure 3:**
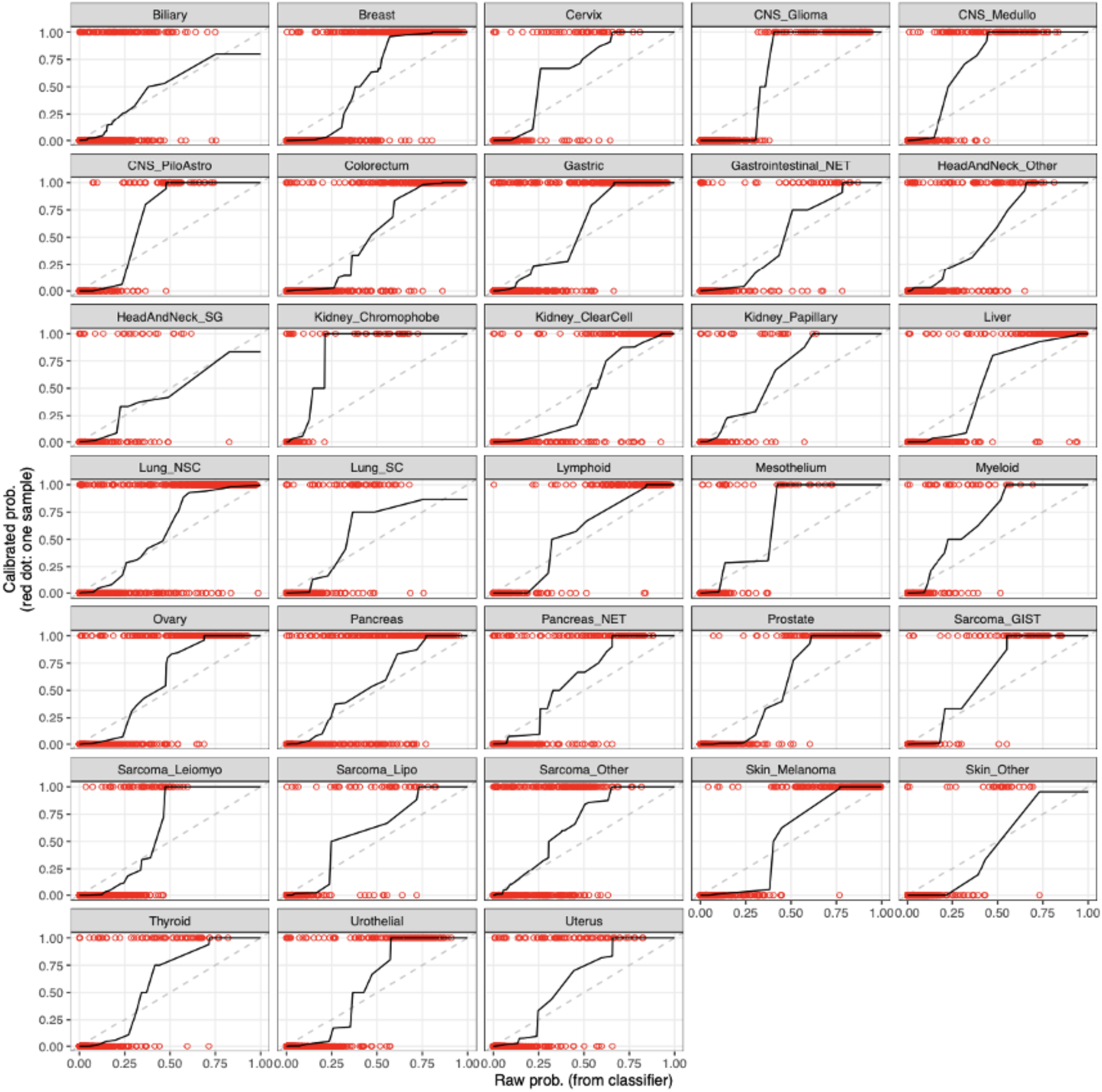
Isotonic regression calibration curves for each random forest in CUPLR. Red dots at y=1 are samples that were predicted by the respective cancer type random forest as that cancer type, whereas dots at y=0 are samples that were predicted as not being that cancer type. Cancer type predictions were obtained by cross-validation.

**Supplementary figure 4:**
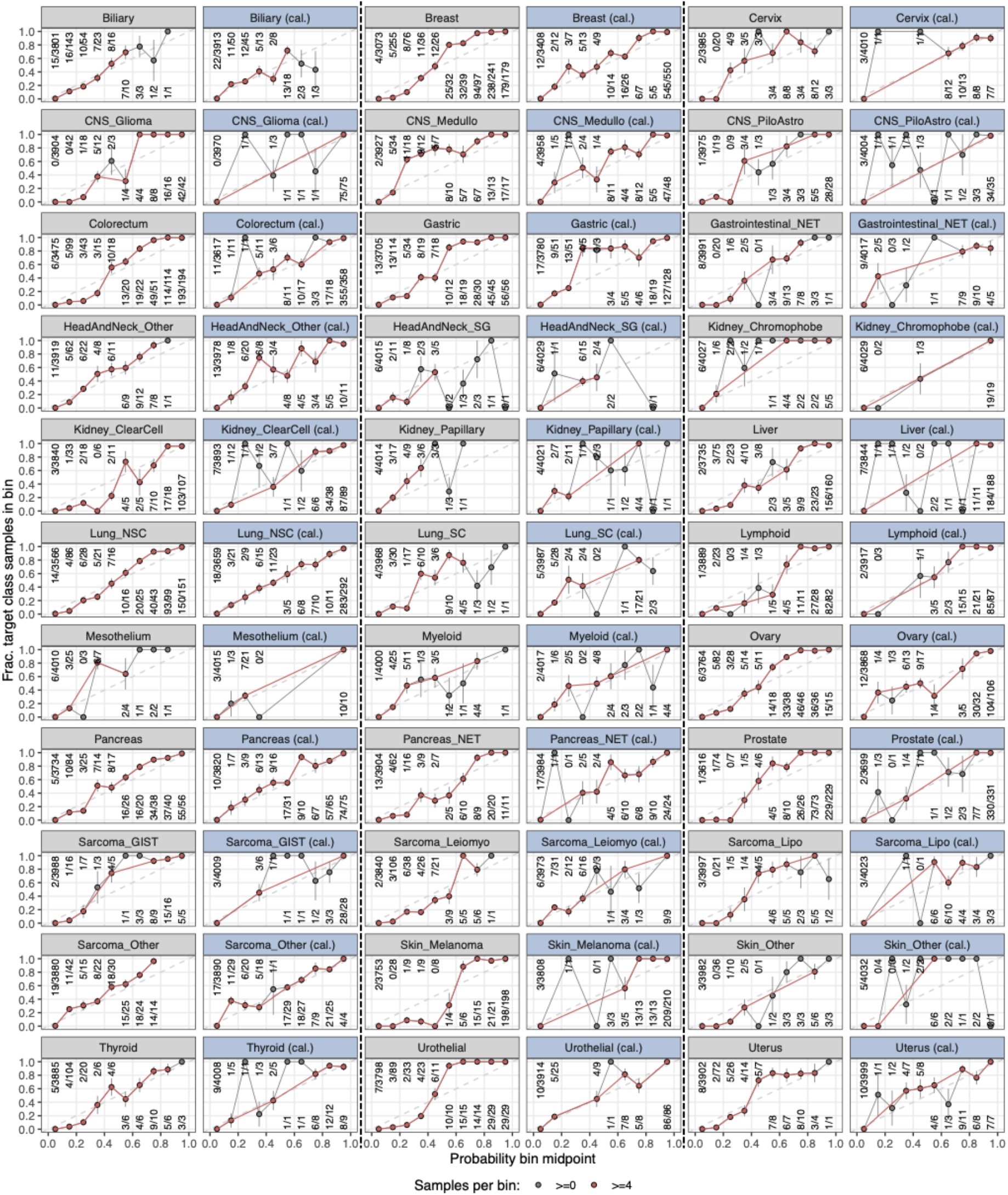
Reliability curves showing the probability biases before and after calibration. Grey panels show the curves before calibration and the blue panels show the curves after calibration. Each dot represents the fraction of samples of the target cancer type in a particular probability bin (a bin at e.g. 0.05 would represent probabilities between 0 and 0.1). Dots above the diagonal are probabilities where the random forest is overconfident whereas dots below the diagonal are probabilities where it is underconfident. A properly calibrated classifier has a reliability curve close to the diagonal. Two curves are shown in each panel, one filtered such that each bin has sufficient (≥4) samples for plotting a stable curve, and one where each bin has ≥0 samples representing the raw reliability curve.

**Supplementary figure 5:**
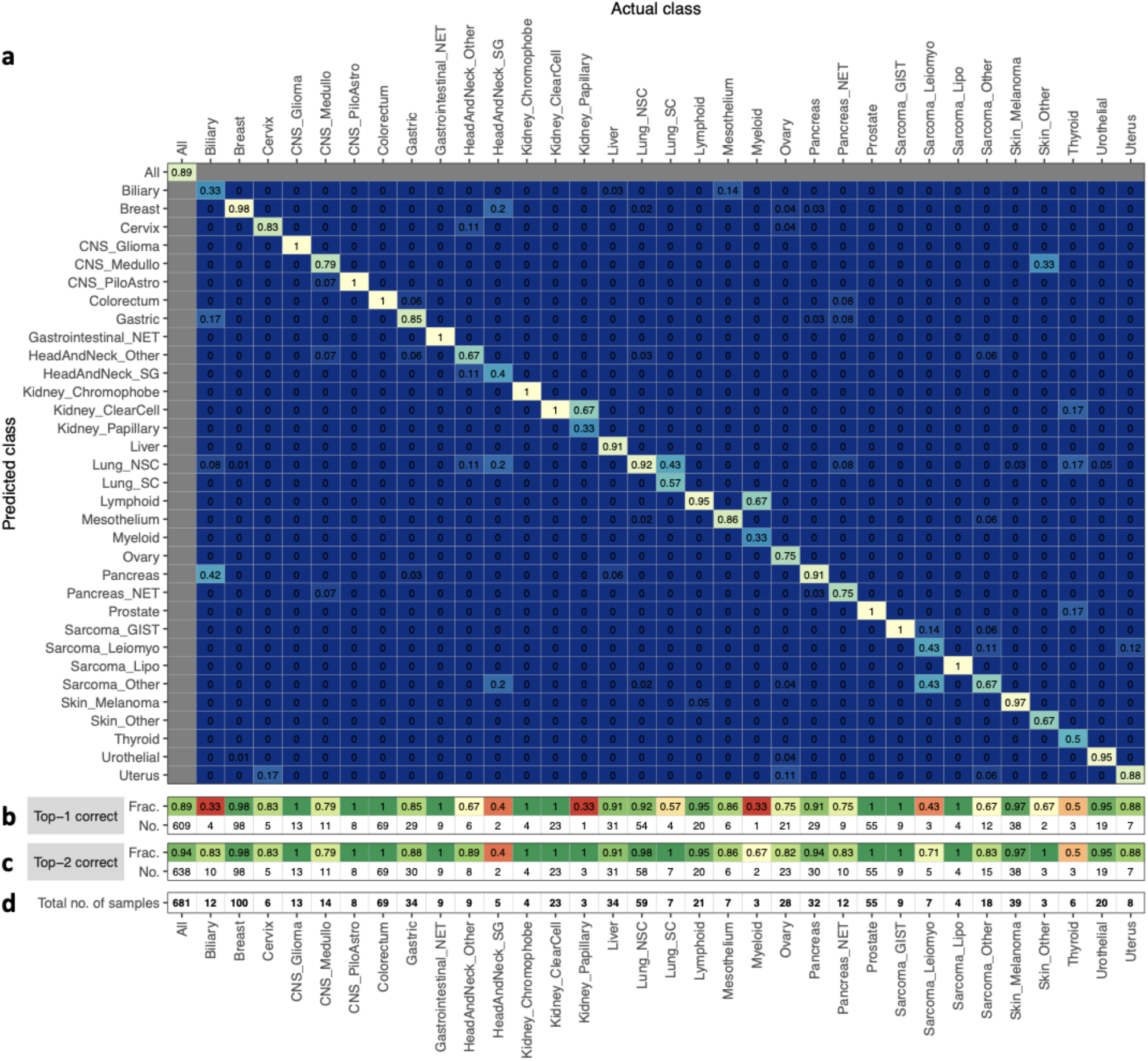
Accuracy of CUPLR based on predictions on the holdout set. **(a)** A heatmap showing the accuracy of CUPLR where columns represent the fraction of samples in a cancer type cohort predicted as a particular cancer type. The diagonal represents the fraction of samples correctly predicted as a particular cancer type (i.e. accuracy). **(b)** The fraction and number of correctly classified samples, equivalent to the diagonal in **(a)**. **(c)** The fraction and number of correctly classified samples when considering the 2 highest probability cancer types. **(d)** The total number of samples for each cancer type. **Abbreviations**; *CNS*: central nervous system; *CNS_Medullo*: medulloblastoma; *CNS_PiloAstro*: pilocytic astrocytoma; *Gastrointestinal_NET*: gastrointestinal neuroendocrine tumor; *HeadAndNeck_SG*: salivary gland cancer; *HeadAndNeck_Other*: head and neck cancers other than salivary gland cancer; *Lung_NSC*: non-small cell lung cancer; *Lung_SC*: small cell lung cancer; *Pancreas_NET*: pancreatic neuroendocrine tumor; *Sarcoma_GIST*: gastrointestinal stromal tumor; *Sarcoma_Leiomyo*: leiomyosarcoma; *Sarcoma_Lipo*: liposarcoma; *Sarcoma_Other*: sarcomas other than leiomyosarcoma, liposarcoma or gastrointestinal stromal tumors; *Skin_Other*: non-melanoma skin cancer.

**Supplementary figure 6:**
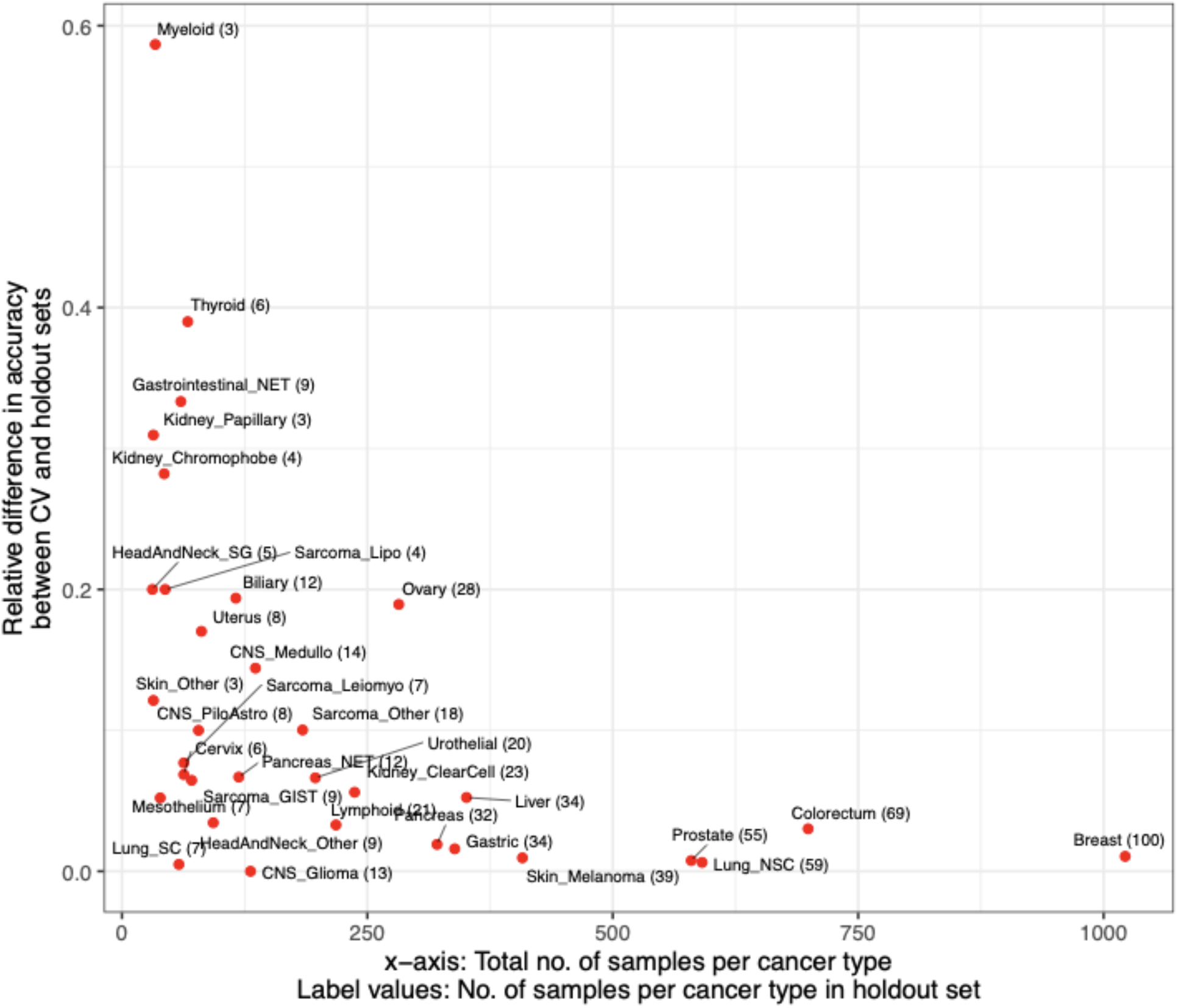
Relative difference in accuracy of CUPLR between the cross-validation (CV) predictions and predictions on the holdout set. The relative difference ranges between 0 and 1 and was calculated using the formula: |a - b| / max(a, b); where a = CV accuracy and b = holdout accuracy. Small cancer type cohorts show higher variation between CV and holdout accuracy.

**Supplementary figure 7:**
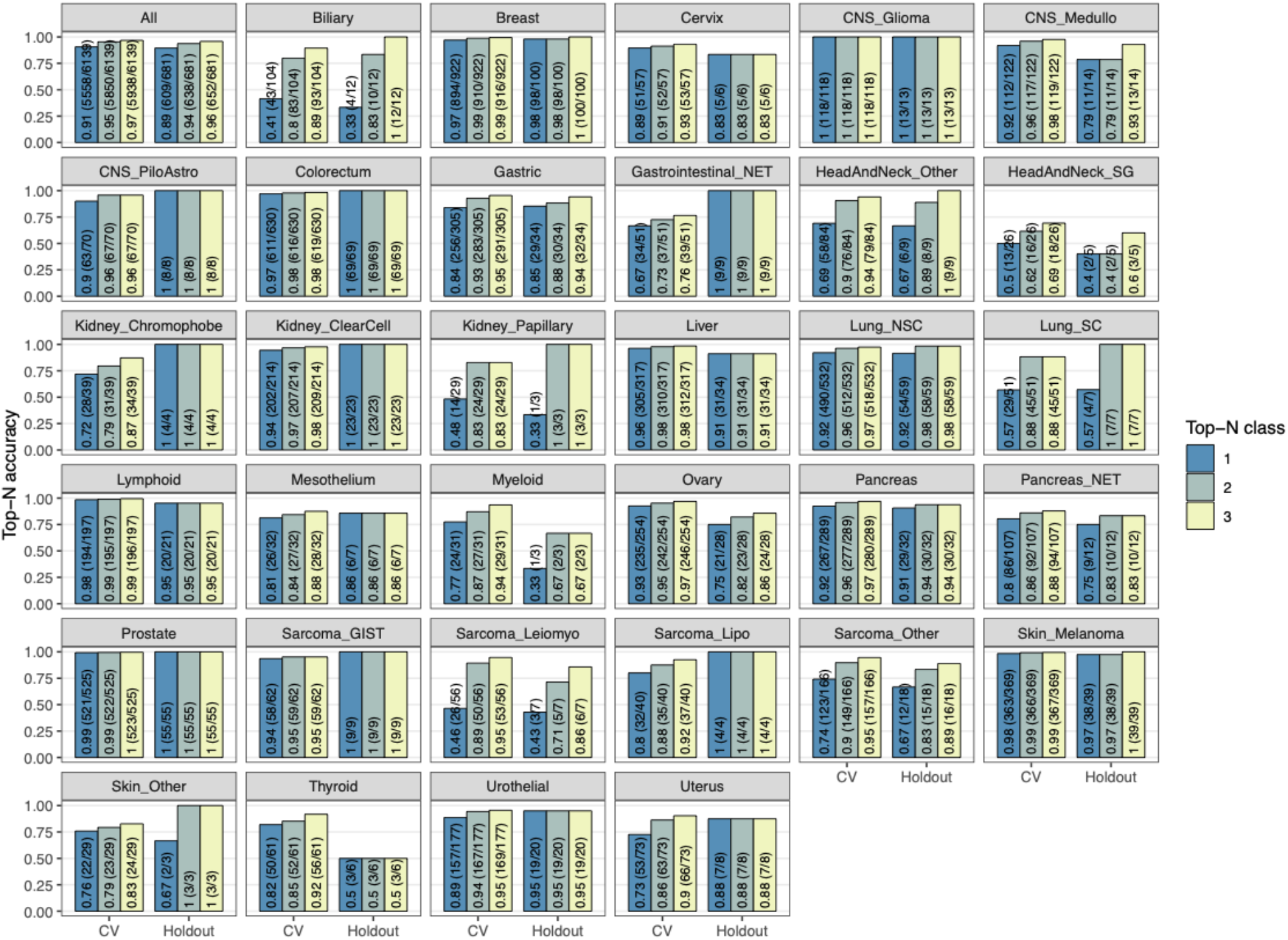
Top-N accuracy of CUPLR based on cross-validation and holdout predictions. Top-3 accuracy would for example represent the fraction of samples for which the correct cancer type is amongst the top 3 predicted cancer types. Bars are labelled with the top-N accuracy value, as well as in brackets the number of correctly predicted samples and the total number of samples.

**Supplementary figure 8:**
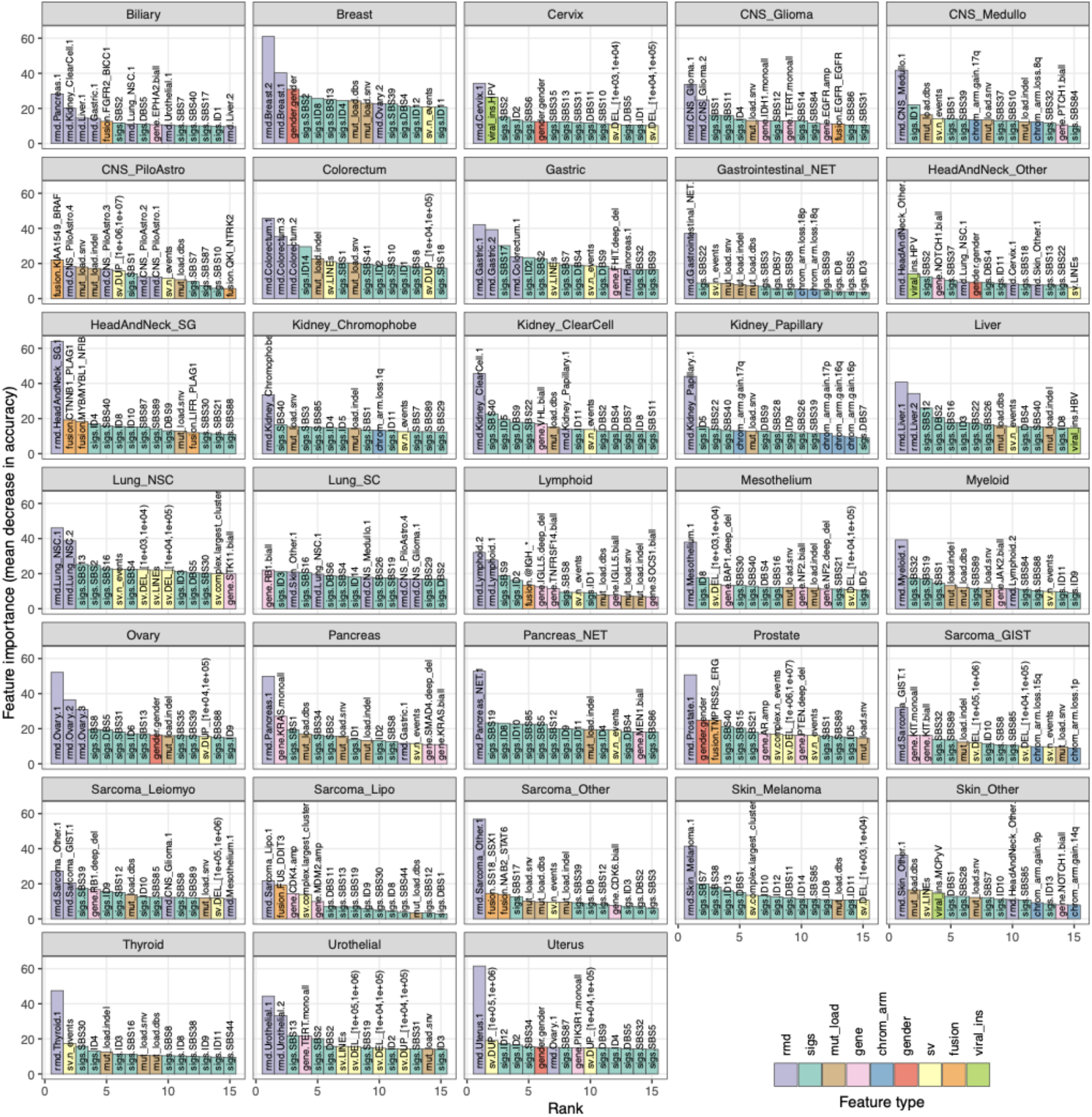
Importance of the top 15 features for each random forest within CUPLR. Feature importance is measured by mean decrease in accuracy across all trees in a random forest upon removing a particular feature. Feature names are in the form {feature type}.{feature name}. **Feature type definitions**; sigs: mutational signatures; rmd: regional mutation density profiles; mut_load: total number of single base substitutions, double base substitutions or indels, sv: structural variants; gene: presence of gene gain or loss of function events; chrom_arm: chromosome arm copy number difference compared to overall genome ploidy; fusion: presence of gene fusions; gender: sample gender as determined by copy number data; viral_ins: presence of viral sequence insertions. For a full description of each feature, see **Supplementary data 3**.

**Supplementary figure 9:**
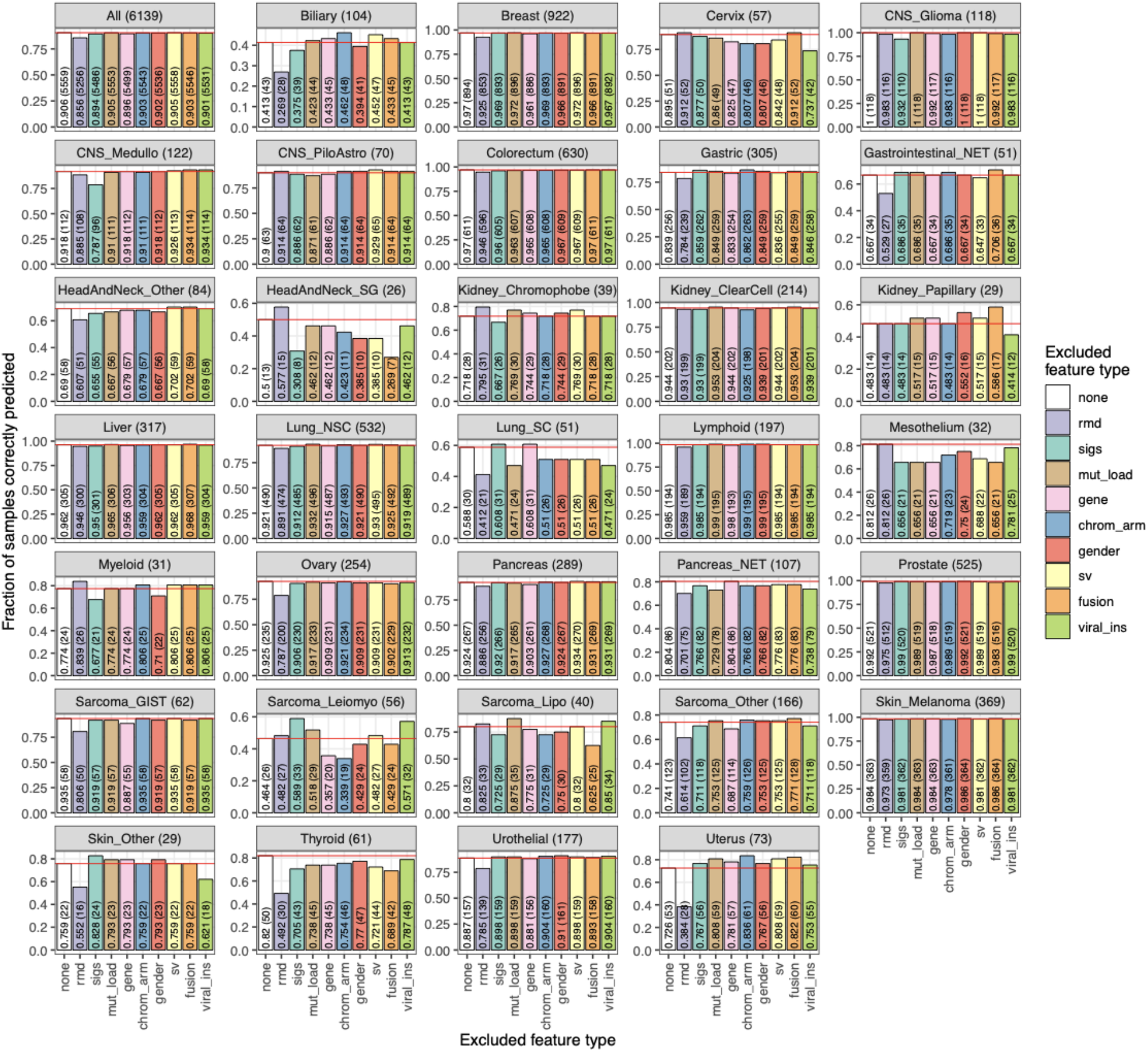
Cancer type prediction accuracy when excluding feature types from the training data. Accuracy (i.e. fraction of sample correctly predicted) was determined using cross validation. Panel titles indicate the cancer type and the total number of samples for that cancer type. Bars are labelled with the accuracy as well as the number of correctly predicted samples. The red line shows the accuracy when no feature types are excluded. **Feature type definitions**; sigs: mutational signatures; rmd: regional mutation density profiles; mut_load: total number of single base substitutions, double base substitutions or indels, sv: structural variants; gene: presence of gene gain or loss of function events; chrom_arm: chromosome arm copy number difference compared to overall genome ploidy; fusion: presence of gene fusions; gender: sample gender as determined by copy number data; viral_ins: presence of viral sequence insertions. For a full description of each feature, see **Supplementary data 3**.

**Supplementary figure 10:**
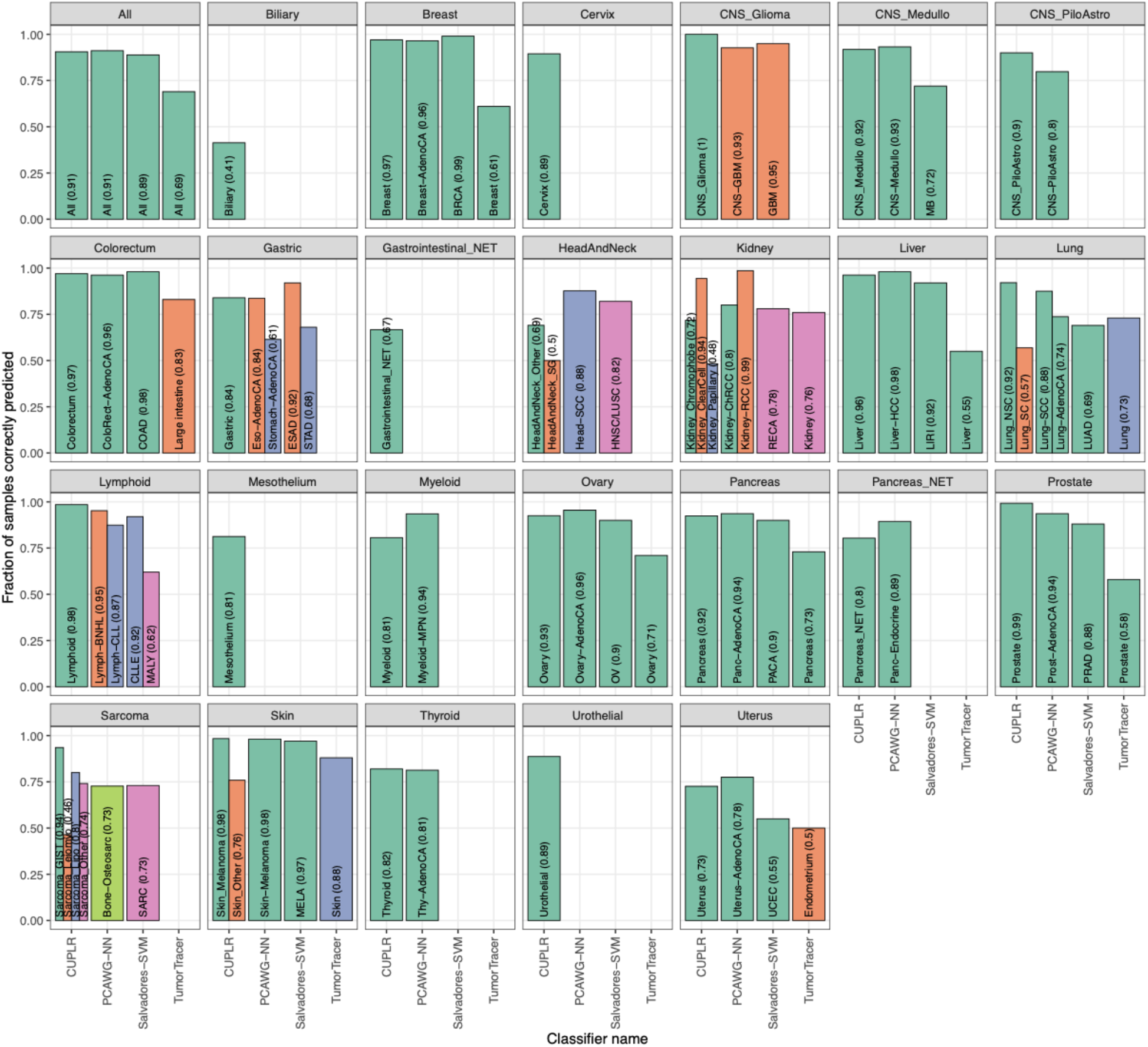
Comparison of cancer type prediction accuracy of CUPLR to other published classifiers. Accuracy (i.e. fraction of sample correctly predicted) for CUPLR was determined using cross validation. Bars are labeled with the cancer type class names from the respective classifiers as well as the accuracy value. Cancer type class names representing the same cancer type across different classifiers are represented by the same bar colors. The names of the published classifiers refer to the following studies; PCAWG-NN: PCAWG neural network by Jiao *et al.* 2020, Salvadores-SVM: support vector machine by Salvadores *et al.* 2019, TumorTracer: Marquard *et al.* 2015.

**Supplementary figure 11:**
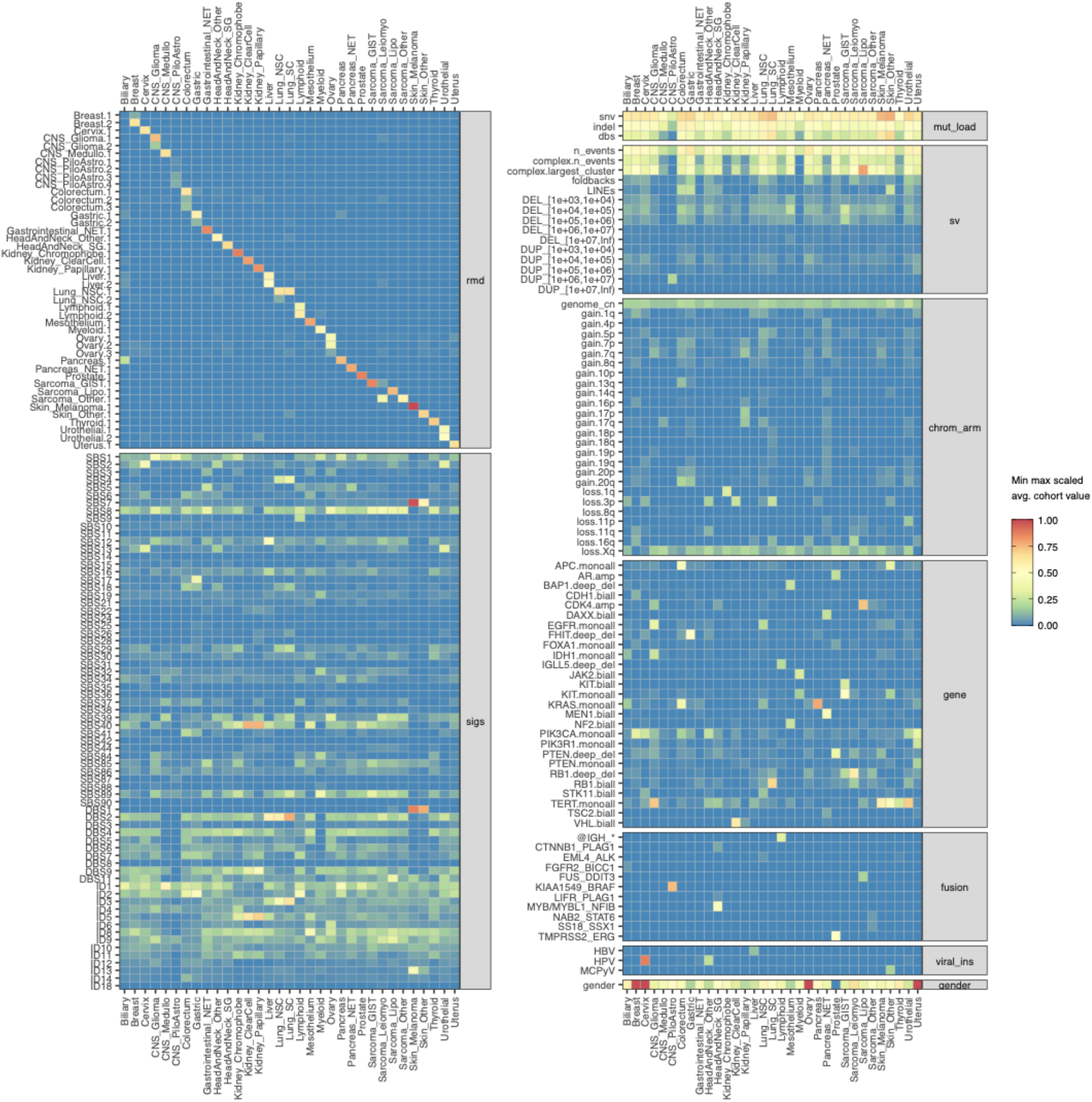
Average scaled feature value per cancer type for the top 200 features (of 496 in total). Feature values are scaled from 0 to 1 based on the minimum and maximum value for each feature across all samples. For gender, feature values towards 1 represent more female samples and values towards 0 more male samples. **Feature type definitions**; sigs: mutational signatures; rmd: regional mutation density profiles; mut_load: total number of single base substitutions, double base substitutions or indels, sv: structural variants; gene: presence of gene gain or loss of function events; chrom_arm: chromosome arm copy number difference compared to overall genome ploidy; fusion: presence of gene fusions; gender: sample gender as determined by copy number data; viral_ins: presence of viral sequence insertions. For a full description of each feature, see **Supplementary data 3**.

### Supplementary notes

#### Impact of confounding factors on performance

Since the RMD profiles and mutational signatures were the most important feature types for CUPLR, we assessed whether the presence of certain confounding factors to these feature types would lead to more incorrect predictions. Firstly, this included DNA repair deficiencies including microsatellite instability (MSI) and homologous recombination deficiency (HRD) which lead to SBS/indel accumulation across the genome and could lead to different RMD profiles than would be expected for a particular cancer type. Secondly, the impact of common chemotherapies including platinum and 5-fluorouracil (5FU) was assessed as treatment induced mutations could also lead to altered RMD profiles. Furthermore, treatment associated mutational signatures could be erroneously predictive of certain cancer types (e.g. platinum signature SBS35 for ovarian cancer) despite them not being intrinsic properties of those cancer types. Lastly, smoking history was assessed for lung cancer patients to determine whether lack of smoking, and by extension the absence of SBS4 (smoking mutational signature), would reduce performance for lung cancer. We found that MSI in gastric cancer and platinum treatment in colorectal cancer did lead to significantly more incorrect predictions (p<0.01, one-sided Fisher’s exact test), though the number of incorrectly predicted samples was low (6 and 4 respectively). However, the presence of the majority of confounding factors did not lead to more incorrect predictions (p≥0.01, one-sided Fisher’s exact test; **Supplementary table 1**).

**Supplementary table 1:** Comparison of the number of correctly and incorrectly predicted samples between patients with and without: microsatellite instability (MSI), homologous recombination deficiency (HRD), smoking history, treatment with platinum, and treatment with 5-fluorouracil (5FU). One-sided Fisher’s exact tests were performed to determine if there were more incorrectly predicted patients with versus without the respective variable. Only cancer types with at least 5 patients being positive for the respective variable (Variable TRUE: Total) are shown.

| Variable | Class | Total | Total with variable data | Variable TRUE |  |  |  | Variable FALSE |  |  |  | p-value |
| --- | --- | --- | --- | --- | --- | --- | --- | --- | --- | --- | --- | --- |
|  |  |  |  | Incorrect | Correct | Total | %Incorrect | Incorrect | Correct | Total | %Incorrect |  |
| has_msi | Gastric | 339 | 339 | 6 | 1 | 7 | 85.7 | 47 | 285 | 332 | 14.5 | 0.00007 |
| has_msi | Breast | 1022 | 1022 | 2 | 8 | 2 | 50.0 | 23 | 989 | 406 | 1.5 | 0.02293 |
| has_msi | Colorectum | 699 | 699 | 2 | 41 | 43 | 4.7 | 19 | 637 | 656 | 2.6 | 0.37481 |
| has_msi | Uterus | 81 | 81 | 3 | 9 | 12 | 25.0 | 13 | 56 | 69 | 26.1 | 0.43682 |
| has_msi | Prostate | 580 | 580 | 0 | 23 | 0 | 0.0 | 4 | 553 | 119 | 20.2 | 1.00000 |
| has_msi | CNS_Glioma | 131 | 131 | 0 | 6 | 0 | 0.0 | 1 | 124 | 67 | 20.9 | 1.00000 |
| has_hrd | Breast | 1022 | 1010 | 8 | 153 | 161 | 6.8 | 13 | 836 | 849 | 1.5 | 0.01115 |
| has_hrd | Sarcoma_Other | 184 | 171 | 4 | 2 | 6 | 66.7 | 40 | 125 | 165 | 24.8 | 0.03874 |
| has_hrd | Pancreas | 321 | 319 | 5 | 26 | 5 | 40.0 | 21 | 267 | 324 | 13.3 | 0.09302 |
| has_hrd | Biliary | 116 | 109 | 6 | 3 | 6 | 16.7 | 56 | 44 | 340 | 4.1 | 0.40053 |
| has_hrd | Lung_NSC | 591 | 583 | 1 | 6 | 1 | 100.0 | 40 | 536 | 59 | 27.1 | 0.40142 |
| has_hrd | Ovary | 282 | 278 | 3 | 105 | 5 | 20.0 | 25 | 145 | 65 | 26.2 | 0.99991 |
| has_hrd | Urothelial | 197 | 192 | 0 | 12 | 108 | 1.9 | 17 | 163 | 170 | 12.9 | 1.00000 |
| has_hrd | Liver | 351 | 346 | 0 | 6 | 1 | 0.0 | 13 | 327 | 233 | 4.7 | 1.00000 |
| has_hrd | Prostate | 580 | 537 | 0 | 55 | 55 | 0.0 | 1 | 481 | 482 | 0.2 | 1.00000 |
| has_smoked | Lung_NSC | 591 | 298 | 11 | 204 | 215 | 5.6 | 10 | 73 | 83 | 13.3 | 0.98792 |
| has_smoked | Lung_SC | 58 | 35 | 16 | 19 | 35 | 45.7 | 0 | 0 | 0 | 0.0 | 1.00000 |
| treated_with_platinum | Colorectum | 699 | 355 | 4 | 40 | 23 | 21.7 | 4 | 307 | 150 | 6.7 | 0.00998 |
| treated_with_platinum | Lung_SC | 58 | 29 | 7 | 2 | 44 | 6.8 | 7 | 13 | 311 | 1.3 | 0.04068 |
| treated_with_platinum | Breast | 1022 | 546 | 2 | 24 | 26 | 7.7 | 7 | 513 | 520 | 1.2 | 0.06391 |
| treated_with_platinum | Lung_NSC | 591 | 173 | 4 | 19 | 2 | 100.0 | 10 | 140 | 16 | 18.8 | 0.09571 |
| treated_with_platinum | Biliary | 116 | 51 | 10 | 10 | 20 | 60.0 | 17 | 14 | 31 | 61.3 | 0.73407 |
| treated_with_platinum | Urothelial | 197 | 96 | 2 | 23 | 8 | 12.5 | 7 | 64 | 15 | 13.3 | 0.73742 |
| treated_with_platinum | Cervix | 63 | 23 | 1 | 7 | 25 | 8.0 | 2 | 13 | 71 | 11.3 | 0.74308 |
| treated_with_platinum | Ovary | 282 | 111 | 4 | 36 | 1 | 100.0 | 9 | 62 | 9 | 77.8 | 0.76261 |
| treated_with_platinum | Gastric | 339 | 231 | 4 | 39 | 40 | 10.0 | 26 | 162 | 71 | 14.1 | 0.85412 |
| treated_with_platinum | Pancreas | 321 | 75 | 0 | 7 | 43 | 9.3 | 8 | 60 | 188 | 13.3 | 1.00000 |
| treated_with_platinum | Prostate | 580 | 342 | 0 | 10 | 0 | 0.0 | 1 | 331 | 2 | 50.0 | 1.00000 |
| treated_with_5FU | Pancreas | 321 | 75 | 2 | 10 | 12 | 16.7 | 6 | 57 | 63 | 9.5 | 0.37691 |
| treated_with_5FU | Colorectum | 699 | 355 | 1 | 26 | 1 | 100.0 | 7 | 321 | 50 | 60.0 | 0.47240 |

As subclonal variants are enriched for treatment induced mutations, we also compared the performance of CUPLR when RMD and mutational signatures were extracted from all mutations versus only clonal mutations (i.e. treatment induced mutations excluded) (**Supplementary figure 12**). No differences in overall accuracy were found (all mutations: 91% vs. clonal mutations 90%), although large differences in accuracy were observed for Gastrointestinal_NET (all mutations: 71% vs. clonal mutations 39%) and Sarcoma_Leiomyo (all mutations: 50% vs. clonal mutations 73%). However, no clear trend of gain or loss in accuracy was observed across the individual cancer types. Treatment induced mutations thus have minimal impact on the performance of CUPLR.

**Supplementary figure 12:**
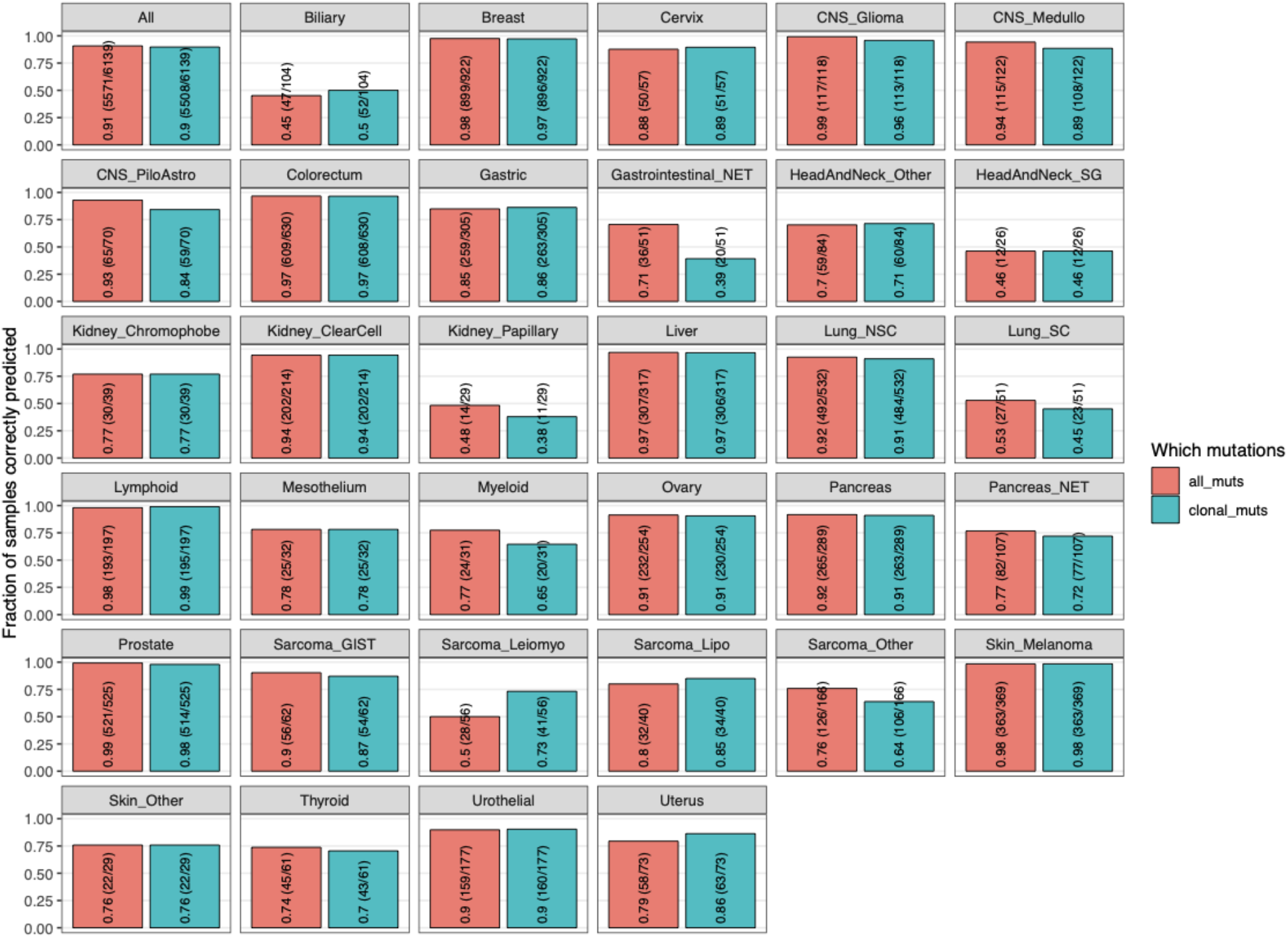
Comparison of cancer type prediction accuracy when using all mutations versus only clonal mutations to generate the regional mutational density and mutational signature features. Accuracy (i.e. fraction of sample correctly predicted) for CUPLR was determined using cross validation. Bars are labelled with the accuracy value, as well as in brackets the number of correctly predicted samples and the total number of samples.

